# Multiple hosts, multiple impacts: the role of vertebrate host diversity in shaping mosquito life history and pathogen transmission

**DOI:** 10.1101/2023.02.10.527988

**Authors:** Amélie Vantaux, Nicolas Moiroux, Kounbobr Roch Dabiré, Anna Cohuet, Thierry Lefèvre

## Abstract

The transmission of malaria parasites from mosquito to human is largely determined by the dietary specialization of *Anopheles* mosquitoes to feed on humans. Few studies have explored the impact of blood meal sources on the fitness of both the parasite and the mosquito. Our study investigated the effects of 3-4 consecutive blood meals from one of four vertebrate species (human, cattle, sheep, or chicken) on several fitness traits, including mosquito feeding rate, blood meal size, susceptibility to wild isolates of *Plasmodium falciparum*, survival, fecundity, F1 offspring development time, and size. Our findings revealed no significant effect on parasite development. Similarly, parasite exposure had no overall effects on mosquito fitness. However, blood meal type did have a strong impact on mosquito feeding rate, survival, lifetime fecundity, and offspring size. Specifically, mosquitoes that were fed successive chicken blood meals produced fewer eggs and fewer and smaller F1 adults compared to those fed human blood. Combining our results in a theoretical model, we show a decrease in the vectorial capacity of mosquitoes fed chicken or cow blood and an increase in the capacity of those fed sheep blood compared to those fed human blood. These findings emphasize the importance of considering the diversity of blood meal sources in understanding mosquito ecology and their role in the transmission intensity of malaria parasites.

## Introduction

The diet of female *Anopheles* mosquitoes, including those species capable of transmitting human malaria parasites, is characterized by the ingestion of vertebrate blood during each ovarian cycle to sustain vitellogenesis and egg production, while plant carbohydrates are mainly used for energy and maintenance reserves (Clements 1992). With a rather short gonotrophic cycle, which can be as fast as 48h between two egg-lays, mosquito females are recurrently looking for a blood meal during which they can transmit malaria parasites. The successful transmission of the malaria parasite is highly dependent of the diet specialization of the *Anopheles* vector and, in particular, its degree of anthropophagy (propensity to feed on human). Female mosquitoes must bite a human host twice to potentially transmit malaria parasites. Therefore, the higher the human feeding rate, the greater the transmission potential (Smith and McKenzie 2004). Furthermore, other key parameters of pathogen transmission such as mosquito longevity (Smith and McKenzie 2004) can be influenced by blood meal source (Lyimo *et al*. 2012, Lyimo *et al*. 2013). Consequently, dietary specialization on human could be associated with fitness benefits for mosquito females which could increase parasite transmission rates.

Comparison of host use among 111 *Anopheline* mosquito populations drawn from 52 species showed that 82% of the populations exhibits some level of dietary specialism (’≥ 50% bloodmeals taken from one host type’; Lyimo and Ferguson 2009). Dietary specialization assumes a trade-off between exploitation of different diets which results in fitness benefits when a specialist feeds on specialized resource and costs when exploiting sub-optimal resources. On the contrary, generalism would be expected when the chances of optimal host encounter are low and the costs of waiting are high (Lyimo and Ferguson 2009) and only small differences between resources would be observed with no optimal use of one type of diet. For example, *Anopheles arabiensis* is rather an opportunistic vector displaying either anthropophilic or zoophilic preferences depending on the geographic area and the relative abundance of humans and cattle (Costantini *et al*. 1999, Takken and Verhulst 2013). On the other hand, in *Anopheles coluzzii*, considered as strongly anthropophagic, environmental changes such as the widespread usage of bed nets can induce mosquitoes to feed on more accessible although less preferred host species (Lefèvre *et al*. 2009). Only a handful of studies have investigated the fitness of anopheline mosquitoes fed on different vertebrate blood (Lyimo *et al*. 2012, Lyimo *et al*. 2012, Lyimo *et al*. 2013, Phasomkusolsil *et al*. 2013, Emami *et al*. 2017). While all studies observed some effects of host type on mosquito fitness traits, the exploitation of less preferred hosts did not seem to strongly impact mosquito fitness so that the predicted relationship between host-preference and fitness benefits was not always confirmed (Lyimo *et al*. 2013) or could be offset by a second blood meal even on a different non –preferred host (Lyimo *et al*. 2012).

Blood meal type is also likely to directly impact parasite fitness since parasite growth is fueled by host resources (Shaw *et al*. 2022). For example, xanthurenic acid, a gametocytogenesis activation factor is synthesized by the mosquito host (Billker *et al*. 1998), and essential amino acids such as valine, histidine and methionine and leucine are incorporated by parasite oocysts (Beier 1998). Similarly, host lipids are taken up by malaria parasites, probably to sustain its membrane biogenesis (Atella *et al*. 2009) while these lipids are also central to mosquito immune defenses and reproduction (Briegel *et al*. 2002, Atella *et al*. 2006, Cheon *et al*. 2006, Rono *et al*. 2010). As vertebrate host species vary in these haematological characteristics (Wintrobe 1933, De Smet 1978, Hawkey *et al*. 1991), they are likely important drivers of both mosquito and parasite fitness. It has also been shown that the provision of a second blood meal to infected females can increase the rates and amount of sporozoites (the mosquito to human infective stage) in salivary glands (Ponnudurai *et al*. 1989, Emami *et al*. 2017, Pathak *et al*. 2022), accelerate parasite growth and shorten the extrinsic incubation period (Brackney *et al*. 2021, Habtewold *et al*. 2021, Kwon *et al*. 2021, Shaw *et al*. 2021, Pathak *et al*. 2022), thereby enhancing the transmission potential of malaria-infected mosquitoes. Thus, the nutritive quality of the mosquito blood meals following malaria parasite invasion might affect parasite fitness, competition for resources between the parasite and its mosquito host as well as mosquito fitness and ability to cope with infection (Shaw *et al*. 2022).

To our knowledge, only two studies have investigated the effects of blood meal sources taken from different vertebrate host species on mosquito competence for malaria parasites by providing a second blood meal 4 or 8 days post-infectious blood meal, using laboratory colonies of mosquitoes, and cultured clones of parasites (Emami *et al*. 2017, Pathak *et al*. 2022). Both studies revealed that the development of the malaria parasite can be influenced by the source of blood consumed following the infection.

While *Anopheles coluzzii* is generally considered highly antropophilic, it can also feed on a wide range of other vertebrate hosts (Lemasson *et al*. 1997, Sousa *et al*. 2001, Caputo *et al*. 2008, Lefèvre *et al*. 2009). The current study investigated the effect of blood meals from four different vertebrate hosts on malaria parasite development and mosquito vector life history traits using field isolates of the parasite *P. falciparum* and a natural population of the mosquito *An. coluzzii* (previously *An. gambiae* M molecular form, (Coetzee *et al*. 2013). Previous studies on females fed on different blood meal sources were carried out after one or two blood meals (Lyimo *et al*. 2012, Lyimo *et al*. 2012, Lyimo *et al*. 2013, Phasomkusolsil *et al*. 2013, Emami *et al*. 2017). In nature, females can be exposed to a wide range of host species and seek a blood meal every 2 to 4 days. Therefore we here used four vertebrate species, provided multiple blood meals (3 to 4) and measured multiple fitness-related traits (feeding rate, blood meal size, competence to parasites, survival, fecundity, F1 development time and wing length) to obtain a thorough picture of the effect of blood-meal diversity on mosquito and parasite fitness. Mosquito females were first fed an infectious or a non-infectious blood meal. They then received up to three subsequent blood meals from either human, chicken, cow or sheep. We predicted that blood type would affect parasite development and mosquito traits such as survival and fecundity and hence vectorial capacity and that effects would add up with subsequent blood meals. Our results were combined into a theoretical model to predict the relative contribution of different vertebrate hosts to overall malaria transmission.

## Methods

### Mosquito colony

Laboratory-reared females of *An. coluzzii* were obtained from an outbred colony established in 2008 and repeatedly replenished with F1 from wild-caught mosquito females collected in Kou Valley (11°23’14”N, 4°24’42”W), 30 km from Bobo Dioulasso, south-western Burkina Faso (West Africa, Fig 1). Females were identified by species diagnostic PCR (Santolamazza *et al*. 2008). Mosquitoes were maintained under standard insectary conditions (27 ± 2°C, 70 ± 5% relative humidity, 12:12 LD). The larvae were reared in spring water under insectary conditions and fed with Tetramin® Baby Fish Food *ad libitum*. Adults were reared in mesh cages (30×30×30cm) and provided with 5% glucose and water on imbibed cottons *ad libitum*. Female mosquitoes were starved for sugar 24h prior access to a blood meal to ensure willingness to feed.

**Figure 1:**
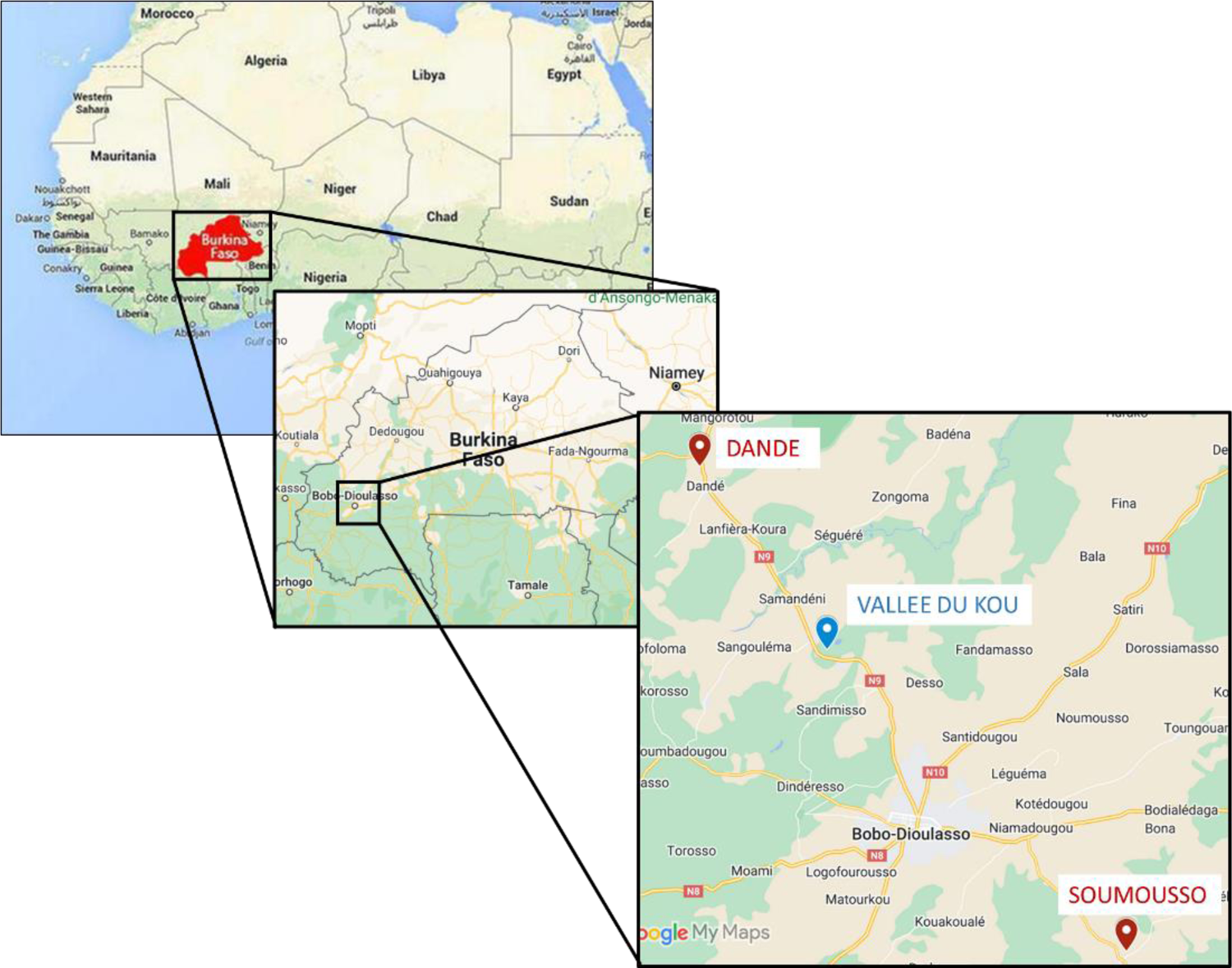
Geographic situation of Burkina Faso in West Africa, and of our sites around Bobo Dioulasso where mosquitoes (Vallée du Kou, blue marker) and parasites (Dande and Soumousso, red markers) were collected.

### Mosquito infection

Experimental infections were carried out as described in (Bousema *et al*. 2012, Ouédraogo *et al*. 2013, Roux *et al*. 2015, Vantaux *et al*. 2015, Vantaux *et al*. 2016). Briefly, 3-to-5-day-old females were fed through membranes on *P. falciparum* gametocyte-infected blood taken from malaria parasite carriers in Burkina Faso. Gametocyte carriers were selected by examining thick blood smears from children aged between 5 and 11 from two villages in southwestern Burkina Faso (Dande and Soumousso, located 60km north and 40km southeast of Bobo-Dioulasso, respectively, Figure 1). Malaria positive individuals were treated according to national recommendations. Venous blood from gametocyte carriers was collected in heparinized tubes. As a negative control (uninfected mosquitoes), females were fed on the same blood in which gametocytes were heat-inactivated. This heat-inactivation inhibits the infection and does not affect the blood nutritive quality (Sangare *et al*. 2013). This was done to avoid the potential confounding effects of different blood origins on fitness of infected and control mosquitoes (Sangare *et al*. 2013, Alout *et al*. 2014, Hien *et al*. 2016, Vantaux *et al*. 2016). Parasite inactivation was performed by placing the blood in a thermo-mixer and heated at 43°C for 15 min and 900 rpm while the remaining blood was maintained at 37°C. Three hundred µl of blood were distributed in membrane feeders maintained at 37°C by water jackets. Cups containing 80 mosquitoes were placed under the feeders to allow blood feeding through Parafilm® membranes for 2 hours. Unfed females were discarded and fed females had access to water only. Five experimental replicates using six distinct parasite isolates were performed (Appendix 1-Table S1, Fig 2). Owing to the high malaria endemicity of Burkina Faso and the resulting high probability of multiplicity of infection (Grignard *et al*. 2018, Sondo *et al*. 2020, Barry *et al*. 2021), although not genotyped we are considering the six isolates as six biological replicates likely having different clonal compositions.

**Figure 2.**
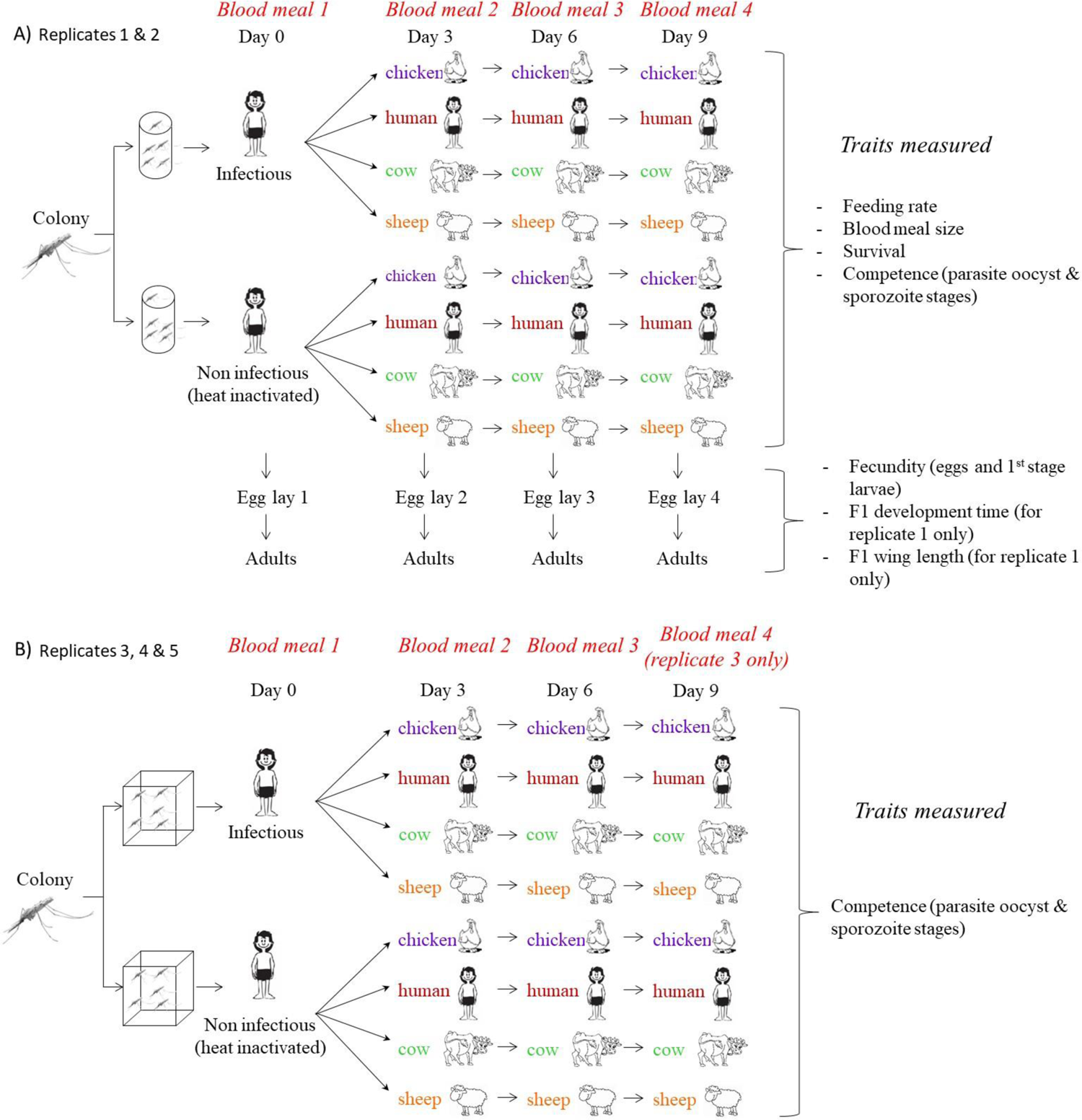
Schematic representation of the experimental design and list of traits measured for the two replicates in cups. (A) and the three replicates in cages (B). For design A, after feeding an infectious blood meal females were distributed in 94 cups and females fed a non infectious blood meal were distributed in 27 cups. Afterwards they received successive blood meals on different vertebrate hosts using membrane feeders. This cup design is schematically represented by cylinders. For design B, after feeding an infectious blood meal or a non infectious blood meal, females were distributed in 32 cages (4 biological replicates by 4 different vertebrates by 2 status of exposure to infection). Afterwards they received successive blood meals on different vertebrate hosts using membrane feeders. This cage design is schematically represented by cubes.

Ethical approval was obtained from the Centre Muraz Institutional Ethics Committee under agreement no. A003-2012/CE-CM. The protocol conforms to the declaration of Helsinki on ethical principles for medical research involving human subjects (version 2002) and informed written consent were obtained from all volunteers.

### Multiple blood meals on different hosts

In addition to the first infectious/uninfectious feed, mosquitoes received two to three additional blood meals every three days through membranes on venous blood drawn from one of four different vertebrate species (“blood type” hereafter): human, cow, sheep or chicken (Fig 2)). After each blood meal, unfed females were discarded and lost to follow-up. As a result, two different experimental designs were employed: one utilizing cups (Fig 2A), which enabled the tracking of small groups of females and the measurement of several life history traits (see below), and a second design using cages (Fig 2B), which allowed for the monitoring of a larger number of females but without measuring all life-history traits. This second design was solely used for measuring mosquito competence and was analyzed separately (see details below). Mosquitoes were fed on the same vertebrate species for either three successive blood meals resulting in a total of four blood meals (replicates 1 to 3) or two successive blood meals resulting in a total of three blood meals (replicates 4 & 5, Fig 2). Membrane feeders were maintained at a specific temperature corresponding to each vertebrate body temperature: 37°C for human blood, 38.5°C for cow blood, 39°C for sheep blood and 41.5°C for chicken blood. For each blood meal episode, three different vertebrate individuals were used per species. The correspondence between mosquito cups and vertebrate individuals was organized so that mosquitoes fed on different individuals of the same host species at each blood meal episode and a total of 14 human volunteers, 8 sheep, 12 cows and 15 chickens were used. Mosquitoes in cages (Fig 2B) were randomly fed on individuals of the same host species at each blood meal and a total of 14 human volunteers, 7 sheep, 8 cows and 18 chickens were used. Fitness costs are more commonly observed in stressful environmental conditions (Lalubin *et al*. 2014, Sangare *et al*. 2014, Roux *et al*. 2015) and sugar feeding strongly affects mosquito survival and fecundity (Foster 2022). Therefore, we did not provide a sugar solution to the mosquitoes during the whole experiment as it could hide or compensate for the fitness effects of the different blood types.

The absence of malaria parasite in human blood donors at feeding episodes 2, 3 and 4 was confirmed by a blood smear prior to blood collection. This study was carried out in strict accordance with the recommendations in the Guide for the Care and Use of Laboratory Animals of the National Institutes of Health. Animals were cared for by trained personnel and veterinarians.

### Traits measurements

The effect of blood type on a series of mosquito life-history traits, namely (i) competence for *P. falciparum* (oocyst / sporozoite prevalence and intensity), (ii) feeding rate, (iii) blood-meal size, (iv) survival, (v) fecundity, (vi) progeny developmental time, and (vii) progeny body size, was assessed.

For replicates 1 and 2, following the first human (infectious or non-infectious) blood meal, a total of 1818 fully-fed *An. coluzzii* females were randomly distributed in 121 paper cups (10 cm height X 7.5 cm down diameter X 9.5 cm up diameter) by group of 9 to 22 (median 12) per cup. Cups were divided in four groups, of which the first three groups of cups contained females fed on infectious blood (thus containing both exposed-infected and exposed-uninfected females) and the fourth group females fed on non-infectious blood. The first (32 cups) and second group (32 cups) were used to investigate the effect of blood type on vector competence, respectively at the parasite oocyst stage 8 days post-bloodmeal (dpbm: days post-bloodmeal) and the sporozoite stage 14 dpbm. The third (30 cups) and fourth group (27 cups) were used to investigate the effect of blood type and *P. falciparum* exposure on mosquito survival. All other life-history traits were measured on mosquitoes from all the cups (n = 121 cups). *F1* traits were measured on offspring from replicate 1. Mosquitoes from replicates 3 to 5, maintained in cages, were used to measure competence traits only (Fig 2).

#### Competence

Oocyst prevalence (i.e. proportion of females harboring at least one oocyst on their midgut) and intensity (i.e. number of *P. falciparum* oocysts in the midgut of infected females) were determined at 8 dpbm. At this time, females had received two additional bloods meal post-infection. Midguts were dissected in 1% Mercurochrome^®^ stain and the presence and number of oocysts were determined under a microscope (at 20× objective). A total of 119 individuals were used to determine oocyst prevalence derived from 13 cups (4 cups from human and sheep blood, 3 cups from cow blood, 2 cups from chicken blood – between 1 to 5 females per cup) in replicate 1 and 16 cups (4 cups per type of blood – between 2 to 11 females per cup) in replicate 2. A total of 757 individuals were used to determine oocyst prevalence in replicates 3 to 5 (202 females in replicate 3, 215 females in replicate 4 and 340 females in replicate 5). Sporozoite prevalence was determined at 14 dpbm. At this time, females had received three (replicates 1 to 3) or two (replicates 4 and 5) additional blood meals post-infection. For each individual, the abdomen was removed and the head and thorax were stored at −20°C. Prevalence was determined using PCR assays on the crushed head and thorax (Morassin *et al*. 2002). Sporozoite prevalence was determined on 34 individuals derived from 6 cups (2 cups from human, cow and sheep) in replicate 1 and 10 cups in replicate 2 (3 cups from human and cow and 4 cups from sheep). Sporozoite prevalence was assessed on 235 individuals in replicates 3 and 5 (175 individuals from replicate 3 and 60 individuals from replicate 5). *Feeding rate*. The feeding rate of females during meals 2 to 4 was calculated as the number of fully-fed females over the total number of females. This trait was assessed on 121 cups for blood meal 2, on 119 cups for blood meal 3 and on 71 cups for blood meal 4. Feeding rate was not measured for the first meal on infectious or non-infectious human blood.

*Mosquito blood meal size* was estimated following each meal (1 to 4) by measuring the amount of haematin (a by-product of the decomposition of haemoglobin) excreted in each paper cup and averaging it by the number of females in the cup (Briegel 1990). One ml of 1% lithium carbonate solution was distributed in each cup to elute faeces and the absorbance of the resulting solution was read at 387nm, using LiCO3 solution as a blank, and compared with a standard curve made with porcine serum haematin (Sigma-Aldrich). Blood meal size was assessed on 116 cups for the first blood meal, on 119 cups for the second blood meal, on 74 cups for the third blood meal and on 60 cups for the fourth blood meal.

Mosquito *survival* was recorded daily at 8:00 by counting and removing the number of dead individuals in the cups. *Survival* was derived from a total of 855 individuals followed from 57 cups.

#### Fecundity

At each gonotrophic cycle (n=4), petri dishes containing humid cotton covered with a piece of Whatmann^®^ paper were placed in the cups two days post-bloodmeal. Eggs laid on the Whatmann^®^ paper were recovered the following morning, pictured and placed in a plastic weighing pan with 25 ml of water. Two days later, pictures of the 1^st^ instar larvae were taken. The number of eggs were counted using the Egg Counter software (Mollahosseini *et al*. 2012) and the number of first instar larvae with ImageJ (Abramoff *et al*. 2004). *Fecundity* was estimated using six parameters: (i) the *egg-laying rate* corresponding to the proportion of cups containing at least one egg, (ii) the *average number of eggs* corresponding to the number of eggs counted in a positive cup (i.e. a cup with at least one egg) divided by the number of females in the cup, (iii) the *lifetime fecundity* corresponding to the sum of the average number of eggs of gonotrophic cycles 2, 3 and 4, (iv) *the hatching rate* corresponding to the number of 1^st^ instar larvae divided by the number of eggs placed in water, (v) the *average number of larvae* corresponding to the number of 1^st^ instar larvae in a plastic weighing pan divided by the number of females in the cup used to collect the eggs placed in that pan, and (vi) the *lifetime production of larvae* corresponding to the sum of the average number of 1^st^ instar larvae of gonotrophic cycles 2, 3 and 4. These six parameters were assessed on a maximum of 121 cups (egglay 1) and a minimum of 41 cups (egglay 4), accounting for mosquito mortality between egglay 1 and 4.

*F1 development time* was assessed by introducing 1^st^ instar larvae (median number of larvae = 9, range = 1-13) randomly selected from each weighing pan in a plastic cup with 50ml of water. Mosquito larvae were provided with Tetramin^®^ Baby Fish Food ad libitum once a day, and excess food was removed to avoid water pollution. Development time was calculated as the duration from egg-lay to emergence and was measured for a total of 1,035 individuals from the four egg-lays of the first replicate (464 individuals from 56 cups in egg-lay 1, 303 individuals from 36 cups in egg-lay 2, 142 individuals from 18 cups in egg-lay 3, 126 individuals from 15 cups in egg-lay 4).

*F1 wing length* was used as a surrogate of body size and was measured from the alula to the wing tip, excluding scales (Van Handel and Day 1989). One wing per F1 individual was dissected on the day following emergence on a subset of individuals. The wing was pictured with a stereomicroscope and measured with ImageJ software (Wayne Rasband, rsb.info.nih.gov/ij/). Wing length was measured on 656 individuals of the first replicate (300 individuals from 55 cups in egg-lay 1, 176 individuals from 34 cups in egg-lay 2, 100 individuals from 18 cups in egg-lay 3, 80 individuals from 15 cups in egg-lay 4).

### Statistical analyses

#### Competence

Parasite prevalence (oocyst or sporozoite stages) and intensity (oocyst stage only) were analysed using Generalized Linear Mixed Models (GLMMs) with a binomial and a zero-truncated negative binomial error structure respectively. The replicates in cups and the replicates in cages were analyzed separately. In these GLMMs, blood type (four levels: cow, sheep, chicken or human blood), gametocytemia and their interaction (only for replicates 3-5) were coded as fixed factors, and cup and parasite isolate nested in replicate (for replicates 3-5) as random factors.

*Feeding rate* was analysed using a GLMM with a binomial error structure. In this model, blood type, *P. falciparum* exposure (two levels: mosquito previously fed an infectious blood meal vs fed the same heat-inactivated blood), blood feeding episode (three levels: 2 to 4) and their interactions as well as parasite isolate were coded as fixed factors and cup as a random factor.

For the following traits, data from the first gonotrophic cycle (resulting from infectious vs. non-infectious human blood) were analysed separately from data from gonotrophic cycles 2-4 for which mosquitoes were fed on four different types of blood (human, cow, sheep, chicken). Data analyses and results from the first gonotrophic cycle are presented in the supplementary material.

#### Mosquito blood meal size

Data from the blood meals 2 to 4 were log-transformed before being analyzed with a GLMM with a Gaussian distribution. In this model, blood type, mosquito exposure, blood-feeding episode and their interactions as well as parasite isolate were coded as fixed factors and cup as a random factor.

*Survival* data were analysed using Cox proportional hazard mixed models (coxme package) with exposure to infectious blood, blood type, parasite isolate and their interactions coded as fixed factors and mosquito cup as a random factor. Since unfed females from blood meals 2 to 4 were removed, they were given a censoring status of 0 indicating that the individual was alive when last seen.

#### Fecundity

Egg-laying rate, hatching rate, average number of eggs (log-transformed), and average number of 1^st^ instar larvae (log-transformed) over gonotrophic cycles 2-4 were analysed using GLMMs with binomial or Gaussian error structures. Blood type, exposure, isolate, blood meal size and gonotrophic cycle were coded as fixed factors and cup as a random factor. In addition, GLMs with quasipoisson structure (to correct for overdispersion) were used to analyze the effect of blood type, exposure, isolate and their interactions on the lifetime fecundity and lifetime production of larvae corresponding to the sum of the average number of eggs and 1^st^ instar larvae over gonotrophic cycles 2-4.

The *development time* of larvae from gonotrophic cycle 2-4 was analyzed using a a Cox proportional hazard mixed effect model with maternal exposure, maternal blood type, gonotrophic cycle, larval density and mosquito sex coded as fixed factors, and rearing cup as a random factor. The effect of blood type on the sex ratio of the progeny was analyzed using a binomial GLMM with blood type coded as a fixed factor and rearing cup as a random factor.

#### F1 wing length

A Gaussian GLMM was used to explore the effects of maternal blood type, maternal exposure, egg-lay episode, larval density and mosquito sex on log-transformed wing length of the progeny from gonotrophic cycles 2 to 4.

For model selection, we used the stepwise removal of terms, followed by likelihood ratio tests (LRT). Term removals that significantly reduced explanatory power (*P*<0.05) were retained in the minimal adequate model (Crawley 2007). All analyses were performed in R v. 3.0.3 (R Core Team 2020). Results are presented as mean ± standard error (se) and proportion ± confidence interval (CI).

### Theoretical modelling

We explored the relative contribution of the blood type on mosquito mean individual vectorial capacity (Saul *et al*. 1990). Individual vectorial capacity (IC) is the mean number of infectious bites given by an infected vector (i.e. the number of bites it gives after the *Plasmodium* extrinsic incubation period is completed). Therefore, IC expresses the efficiency with which individual mosquitoes transmit malaria. To estimate IC, we developed a model that simulates the daily life history of individual mosquito vectors after taking an infectious blood meal on a human under various scenarios (Fig 3). The environment (= scenario) was characterized by the presence of humans and an alternative host (either chicken, cow or sheep) with varying availability (0 to 3 consecutive possible feeding attempts during *Plasmodium* incubation period). There was therefore 12 scenarios tested (3 alt. host x 4 availability levels). 250 000 individuals (representing 500 populations of 500 individuals) were simulated per scenario. The model allowed to track daily physiological states (either Host-Seeking, HS; Blood-Fed, BF; or Resting, R) of individuals. Daily transitions from one state to another depended on survival probability (related to the origin of the previous blood meal) and blood-feeding success probability (related to the host that the mosquito is attempting to bite: human, chicken, cow or sheep; only for transition from HS to BF). A binomial GLMM of feeding success and a COXPH model of survival were fitted to the data presented in the manuscript and used to calculate host-specific probabilities of feeding success and daily survival. For each individual simulation, the number of days spent in state BF (= number of successful feeding attempts) following the duration of *Plasmodium* extrinsic incubation period (n = 11 days) was counted and the mean (= IC) was calculated for each population. The model was implemented in R with the use of the tydiverse, furr, glmmTMB, coxme and emmeans packages (Brooks *et al*. 2017, Wickham *et al*. 2019, Therneau 2020, Vaughan and Dancho 2021, Lenth 2022). The detailed description of the model following the ODD (overview, design concepts and details) protocol for describing individual- and agent-based models (Grimm *et al*. 2010) is as follow:

**Figure 3:**
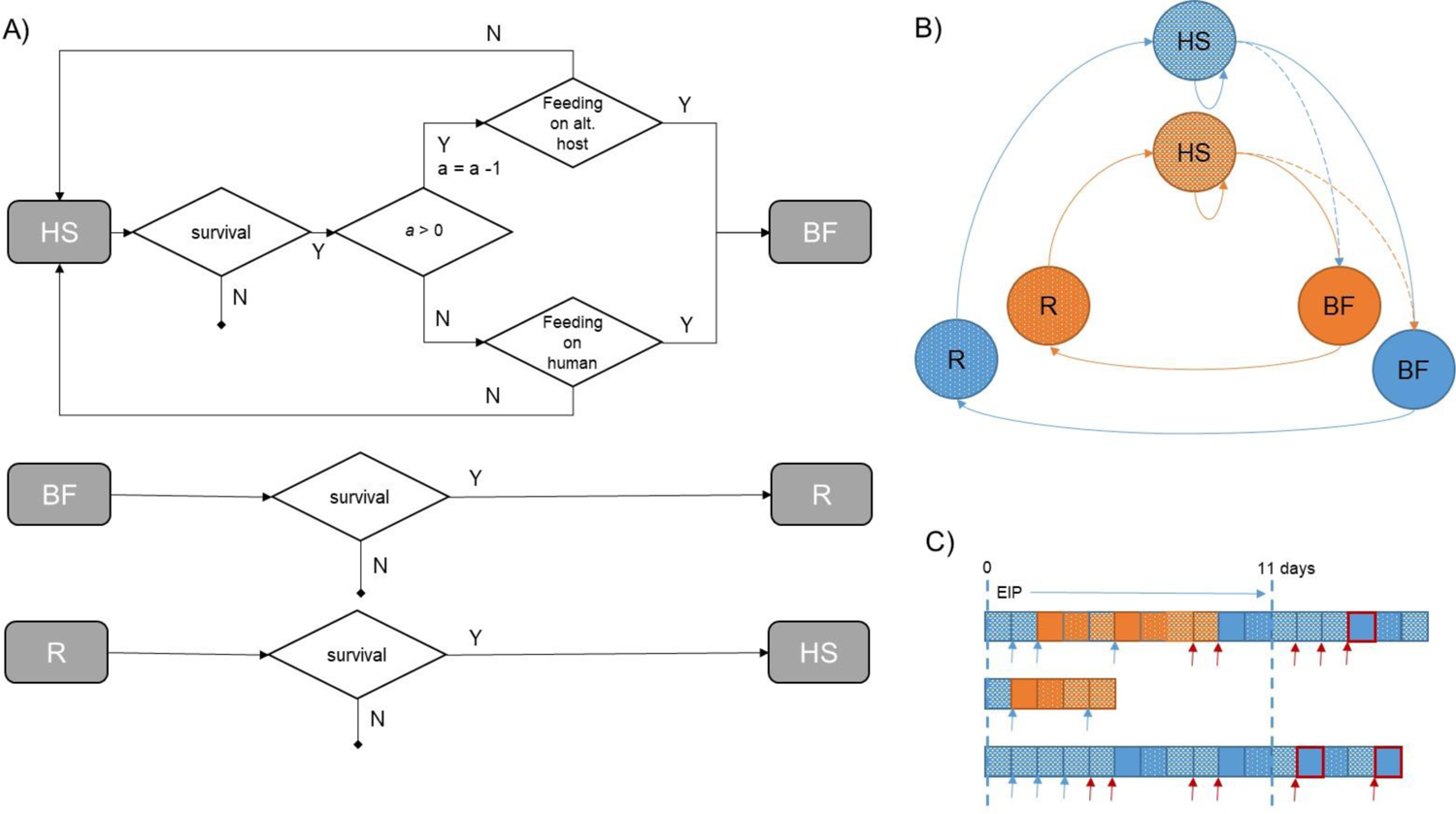
Schematic representations of the model and its outputs. (A) Daily transitions between physiological states (HS: Host-Seeking; BF: Blood-Fed; R: Resting) and the process which cause these transitions (Y: Yes; N: No; alt: Alternative; *a*: remaining number of feeding attempts to be done on the alternative host). (B) Diagram showing possible successions between physiological states together with the origin of blood-meal (blue: human blood, orange: alternative host). At the start of a simulation, the mosquito is always HS with a previous blood meal taken on human (blue). If *a* > 0, it attempts to feed on an alternative host. If successful (blue doted arrow), it enters the orange cycle where daily survival probabilities depends on the type of alternative host bitten, until *a* = 0. Then, the mosquito attempts to feed on human and if it succeeds (orange doted arrow), it enters the blue cycle, with corresponding daily survival probabilities, until death. (C) Three possible sequences of physiological states for single mosquitoes under a scenario of up to 3 possible feeding attempts on an alternative host, until death. Each box corresponds to one day. Colors and texture of boxes relates to panel B. Blue and red arrows indicate feeding attempts on alt. host and human, respectively. Boxes with red borders indicate days in state BF following EIP (External Incubation Period) of the malaria parasite, that are counted to compute Individual Vectorial Capacity (IC).

#### Purpose

The purpose of the model is to explore the effect on the *Anopheles* mean individual vectorial capacity (1) of various number of feeding attempts on alternative animal hosts during the *Plasmodium* extrinsic incubation period.

#### Entities, state variables and scales

The entity of the model is a female *Anopheles coluzzi.* having taken an infectious blood meal on a human. The female *Anopheles* is characterized by its physiological state (Host-seeking, HS; Blood fed, BF; or Resting, R), the source of its last blood meal (human or animal) and the remaining number of feeding attempts to be done on the alternative host. The environment is characterized by the presence of humans and an alternative host (either chicken, cow or sheep) with varied availability (x consecutive possible feeding attempts). One time step of the model corresponds to one day. Simulation is run for 40 days or until the female dies. Every simulation (individual *Anopheles*) is independent (i.e. no interaction).

#### Process overview and scheduling

Every time step, the physiological state of the female mosquito is updated according to its state at the previous time step, survival probability (depending on the origin of the previous blood meal) and blood-feeding success probability (depending on the host that the mosquitoes is attempting to bite: human, chicken, cow or sheep; only for transition from HS to BF). An HS female, if survives, attempts to feed and becomes BF (if it successes to feed) or stay HS (if it fails to feed) at next time step. A BF female, if survives, becomes R. An R female, if survives, becomes HS. Firsts feeding attempts are made on the alternative animal host (the total number of possible attempts is defined by the scenario: 0, 1, 2 or 3). Then, all attempts are made on humans. The sequence of successive daily physiological states is stored.

#### Design concept

Mosquito-to-human transmission may occur during each bite after the pathogen incubation period is completed. Transmission is therefore dependent on both the longevity of the mosquito and the duration of its gonotrophic cycle (i.e. the mean time between two consecutive bites). Longevity vary according to the daily survival probability, duration of the gonotrophic cycle vary according to feeding success probability (as feeding is delayed in case of failure).

*Stochasticity* occurs for daily transition between physiological states since feeding success and survival are Bernoulli trials with host-specific probabilities of success calculated from (i) a binomial GLMM of feeding success and (ii) a Cox proportional hazard model of survival (see sub-model section below), respectively.

#### Observation

For each individual simulation, the number of days spent in state BF (= number of successful feeding attempts) following the duration of *Plasmodium* extrinsic incubation period (n = 11 days) is counted and population mean is calculated. Populations are made of 500 independent individuals.

#### Initialization

Initially, a female vector is in physiological state “Blood-Fed” with previous blood meal taken on human (the infectious blood meal).

#### Sub-models

Probabilities used in the Bernoulli trial of feeding success were provided by a binomial mixed effect model of feeding success fitted on data from the current study. Feeding success was modelled according to blood meal source and with blood meal episodes and cup of origin (of the mosquito batch) as crossed random effect, using the *glmmTMB* function. Feeding success probabilities according to blood meal source were extracted using the *emmeans* function and are shown in Table 1. Probabilities used in the Bernoulli trial of survival were derived from the result of a Cox Proportional Hazard mixed effect model of survival fitted on data from the current study. Death events were modelled according to blood meal source and replicates with cups of origin (of the mosquito batch) as random effect, using the *coxme* function. Hazard ratios (relative to human blood source) were extracted using the *emmeans* function (Table 1). In the simulations, we set daily survival probability with a previous blood meal taken on human to be 0.8. Survival probabilities for mosquitoes having taken their previous blood meal on animals was therefore 0.8 exponentiated by the corresponding Hazard ratio (Table 1).

**Table 1:** Hazard ratio, corresponding daily survival probabilities, and feeding success probabilities used in the individual simulations.

| Host | Hazard Ratio | Daily survival probability | Feeding success probability |
| --- | --- | --- | --- |
| Human | 1 (ref) | 0.8 | 0.753 |
| Chicken | 3.254 | 0.484 | 0.796 |
| Cow | 1.392 | 0.733 | 0.717 |
| Sheep | 0.833 | 0.83 | 0.774 |

## Results

### Competence

In replicates 1 and 2, among the 119 females dissected eight days post-infectious blood meal, 71 (59.7 ± 9 %) harboured parasites. Parasite prevalence, intensity and gametocytemias for each parasite isolate are given in the supplementary material (Appendix 1-Table S1). Although blood type did not significantly influence oocyst prevalence (*X*^2^3 = 5.02, P = 0.17; Fig 4A), oocyst intensity varied among blood type (*X*^2^3 = 10.55, P = 0.01; Fig 4A). However, none of the multiple post-hoc comparisons were significant. As expected, the parasite isolate with the highest gametocytemia (A: 232 gametocyte/µl) caused higher parasite prevalence and intensity than that with the lowest gametocytemia (32 gametocyte/µl) (prevalence: 77.8 ± 14% and 51.8 ± 11%, *X*^2^1 = 6.92, P = 0.009; intensity: 21.1 ± 3.6 oocysts and 6.7 ± 0.7 oocysts; *X*^2^1 = 24.89, P < 0.0001). In replicates 1 and 2, prevalence at the sporozoite stage was determined in individuals fed on cow, human and sheep only since all the females fed on chicken blood were dead by 14 dpi. Blood type and parasite isolate did not influence sporozoite prevalence (*X*^2^2 = 0.62, P = 0.73 and *X*^2^1 = 2.09, P = 0.15, respectively). The prevalence at the oocyst and sporozoite stages were similar for both isolate A (77.8 ± 13% *vs*. 90 ± 19%, respectively; Fisher’s exact test: n=46, P = 0.66) and B (51.8 ± 11% *vs*. 66.7 ± 19%, respectively; n=107, Chi square test: *X*^2^1 = 1.12, P = 0.29).

**Figure 4.**
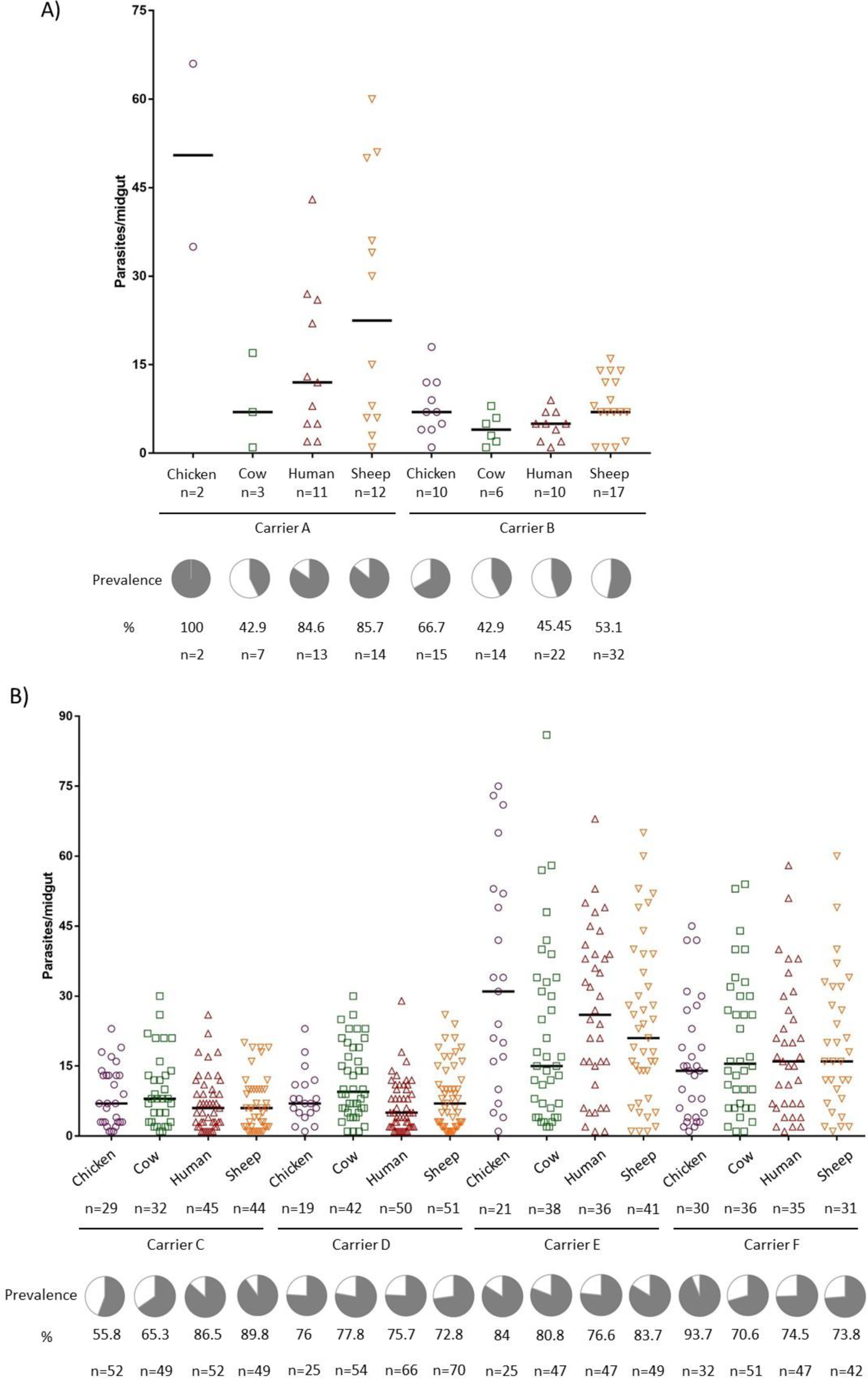
Effects of blood type on parasite (oocyst stage) prevalence and intensity for each parasite isolate in (A) replicates 1 (isolate A) & 2 (isolate B) and in (B) replicates 3 (isolate C), 4 (isolate D) & 5 (isolates E and F). Horizontal bars show the median values. Each colored point represents a *P. falciparum*-infected mosquito individual. Pies show the infection prevalence (grey area). Numbers indicate the sample size (n = total number of mosquito females for parasite prevalence or number of infected females for parasite counting) for each treatment and isolate.

In replicates 3 to 5, among the 757 females dissected eight days post-infectious blood meal, 580 (76.6 ± 3%) harboured oocysts (Appendix 1-Table S1). Neither blood type (*X*^2^3 = 3.14, P = 0.37) nor gametocytemia (*X*^2^1= 1.37, P = 0.24), nor their interaction (*X*^2^3= 1.27, P = 0.74) affected oocyst prevalence (Fig 4B). Parasite intensity did not significantly vary among blood type (*X*^2^3= 4.99, P = 0.17; Fig 4B). Gametocytemia was positively correlated to parasite intensity (*X*^2^1=11.09, P < 0.001) and there was a significant blood type by gametocytemia interaction (*X*^2^3= 11.06, P = 0.01). Parasite prevalence at the sporozoite stage was not significantly affected by the blood type (*X*^2^2 = 0.65, P = 0.72), nor by gametocytemia (*X*^2^1 = 1.3, P = 0.25) nor by their interaction (*X*^2^2 = 0.33, P = 0.85). Parasite prevalence at the oocyst and sporozoite stages were similar for isolate C (74.3 ± 6% *vs*. 81.7 ± 6% respectively; n=377, Chi square test: *X*^2^1 = 2.6, P =0.11), E (80.9 ± 6 % *vs*. 71.4 ±14 %, respectively; n=210, Chi square test: *X*^2^1 = 1.31, P =0.25), and F (76.7 ± 6% *vs*. 66.7 ±22 %, respectively; Fisher’s exact test: n=190, P = 0.39).

### Feeding rate

Blood type significantly affected mosquito feeding rate (X^2^3 = 14.4, P = 0.002) with highest overall feeding on sheep blood (66.32±1.8%) followed by chicken blood (64.6±1.9%), human blood (64.1±1.9%) and cow blood (58.8±1.9%). There was a significant interaction between blood type and feeding episode (X^2^6 = 37.8, P< 0.0001; Fig 5A), with cow blood providing lowest feeding rate during the second blood-meal and highest rate during blood-meals three and four. Mosquitoes that received an infectious blood meal displayed lower feeding rate than mosquitoes previously fed an uninfectious blood-meal (61.4 ± 1.9% *vs*. 71.1 ± 1.8%; X^2^1 = 9.09, P = 0.003, Fig 5B), regardless of the blood type (exposure: blood type: X^2^3 = 0.38, P=0.94, Fig 5B) and of the feeding episode (exposure:feeding episode: X^2^2 = 0.03, P=0.99). Feeding rate consistently increased over the successive feeding episodes (53.1 ± 2, 81.2 ± 1.5, and 89.1 ± 1.2% at the 2^nd^, 3^rd^ and 4^th^ episode, respectively; X^2^2 = 160.2, P< 0.0001, Fig 5A). No other effects were found (Appendix 2-Table S2).

**Figure 5.**
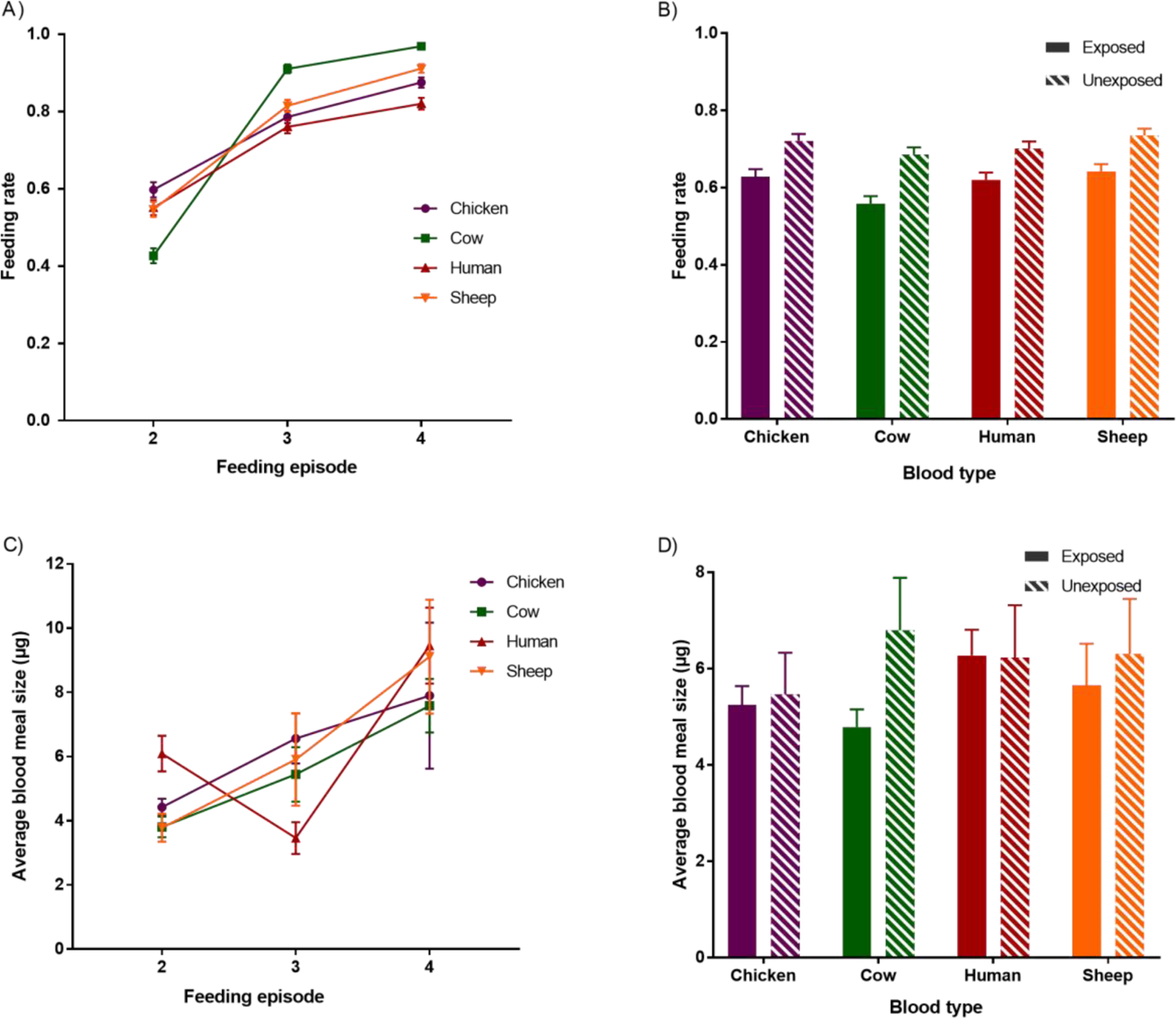
Effects of blood type on mosquito feeding rate and blood meal size. (A) Feeding rate (number of fed females/number of alive females) ± 95% CI as a function of blood feeding episode and blood meal type. B) Feeding rate as a function of blood meal type and infection group (females exposed vs. females unexposed to an infectious blood meal on feeding episode 1). Bars show the average feeding rate across feeding episodes 2 to 4 + 95% CI. C) Average blood meal size ± se as a function of blood feeding episodes and blood meal type. D) Average blood meal size + se as a function of blood type and infection group (females exposed vs. females unexposed to an infectious blood meal). Bars show the average meal size across feeding episodes 2 to 4.

### Mosquito blood meal size

Blood type did not significantly affect meal size (*X*^2^3 = 4.2, P = 0.24, Fig 5C). Meal size varied among feeding episodes (*X*^2^2 = 54.04, P < 0.0001) with biggest size observed for the fourth bloodmeal. There also was a significant blood type by feeding episode interaction (*X*^2^6 = 23.7, P = 0.0006; Fig. 5C) such that blood type providing highest or lowest meal size were not always the same across feeding episodes. Meal size was not influenced by the previous exposure of mosquitoes to parasites (*X*^2^1 = 0.18, P = 0.67) regardless of the blood type (exposure: blood type: *X*^2^3 = 4.7, P = 0.19; Fig 5D) or the feeding episode (exposure: feeding episode: *X*^2^2 = 0.8, P = 0.67). No other effects were found (Appendix 2-Table S2).

### Survival

Mosquito survival over the duration of the experiment was strongly influenced by the blood type (X^2^3 = 68.26, P < 0.0001; Fig 6), with lowest survival observed when females were successively fed with chicken blood (median survival time: chicken: 6 days, cow: 9 days, human: 10 days and sheep 11 days). Females fed on isolate B during the first feeding episode survived significantly longer than females fed on isolate A (10 and 6 days respectively; X^2^1 = 24.38, P < 0.0001, Appendix 3-Fig S1). Mosquito exposure to *P. falciparum* gametocytes did not significantly influence mosquito survival (9 days for both unexposed and exposed mosquitoes, X^2^1 = 0.03, P =0.87) regardless of the blood type (exposure: blood type: X^2^3 = 2.38, P=0.50; Fig 6) or the isolate (exposure: isolate: X^2^ =0.27, P =0.96). No other effects were found (Appendix 2-Table S2).

**Figure 6.**
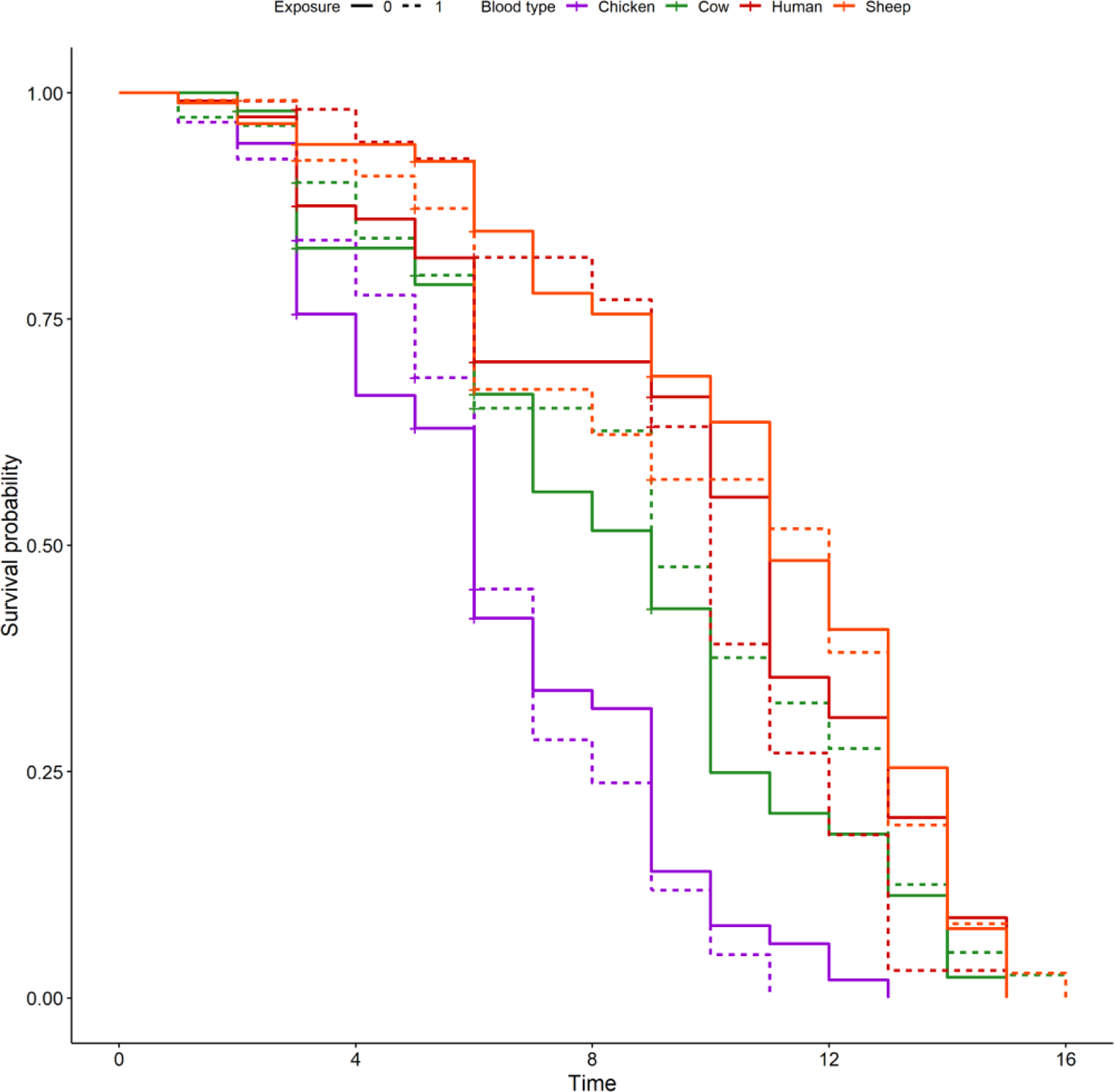
Effects of blood type (colored lines) and infection group (dashed lines: females exposed vs. solid lines: females unexposed to an infectious blood meal on feeding episode 1) on mosquito survivorship.

### Fecundity

Blood type had no effect on egg-laying rate (X^2^3 = 5.6, P = 0.13; Fig 7A), the average number of eggs per female (X^2^3 = 0.85, P = 0.84; Appendix 3-Fig S2A), the hatching rate (X^2^3 = 5.26, P = 0.15; Fig 7B), nor on the average number of 1^st^ instar larvae (X^2^3 = 1.4, P = 0.70; Appendix 3-Fig S2B).

**Figure 7.**
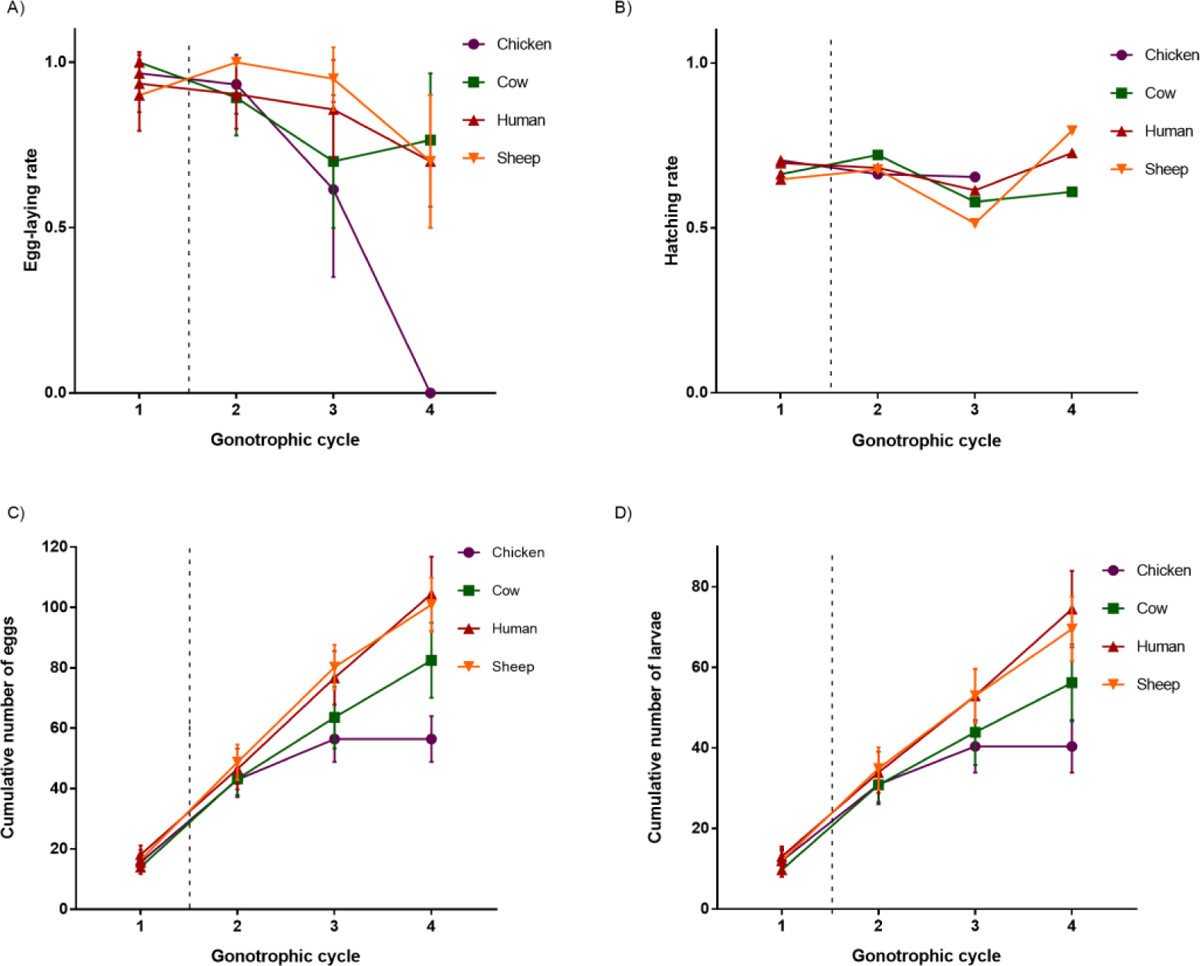
Effects of blood type on mosquito fecundity and fertility. (A) Egg-laying rate (number of cups with eggs/total number of cups) ± 95% CI as a function of gonotrophic cycles and blood type. B) Hatching rate (number of larvae/number of eggs) ± 95% CI as a function of gonotrophic cycles and blood type. C) Cumulative average number of eggs ± se as a function of gonotrophic cycle and blood type. D) Cumulative average number of larvae ± se as a function of gonotrophic cycles and blood type.

Importantly, blood type significantly influenced mosquito lifetime fecundity i.e. the cumulative average number of eggs at the 4^th^ gonotrophic cycle (F1,88=4.79, P=0.004; Fig 7C as well as the lifetime production of larvae i.e. the cumulative average number of larvae at the 4^th^ gonotrophic cycle (F1,88=3.86, P=0.012; Fig 7D), with females successively fed on chicken having a lower lifetime fecundity and lifetime production of larvae than females fed on human blood (z=3.29, P=0.005 and z=3.03, P=0.01) or on sheep blood (z=3.17, P=0.008 and z=2.72, P=0.03). All other post-hoc comparisons were not significant. The effect of blood type on lifetime fecundity and on lifetime production of larvae was independent of the isolate used during the first feeding episode (blood type: parasite isolate: F1,76=0.58, P=0.63 and F1,80=0.31, P=0.82) and of exposure (blood type: exposure: F1,80=0.67, P=0.57 and F1,80=0.86, P=0.46).

Parasite exposure did not influence egg-laying rate (X^2^1 = 0.08, P = 0.78), the average number of eggs (X^2^1 = 0.0005, P = 0.98; Appendix 3-Fig S2C), the lifetime fecundity (F1,84=0.005, P=0.94), the hatching rate (X^2^1 = 2.06, P = 0.15), the average number of 1^st^ instar larvae (X^2^1 = 0.97, P = 0.32, Appendix 3-Fig S2D) nor the lifetime production of larvae (F1,84=0.07, P=0.79).

The successive gonotrophic cycles influenced the egg-laying rate (X^2^2 = 53.6, P < 0.0001; Fig 7A) with 93.3± 4% of egg-positive cups following egg-lay 2, 79.7±9% following egg-lay 3 and 68.3±12% following egg-lay 4. This reduction was likely associated to mosquito mortality causing decreased mosquito number in the cups overtime. Indeed, over the four gonotrophic cycles the probability of laying eggs was positively associated to the survival rate (X^2^1 = 32.9, P < 0.0001; Appendix 4-Supplementary methods and results; Appendix 3-Fig S3A). This was particularly true for the chicken blood treatment for which there were only 3 cups left at cycle 4. Similarly, the hatching rate varied over the successive gonotrophic cycles (X^2^2 = 501, P < 0.0001) with maximum rate observed for egg-lay 4 (71.7 ± 0.5 %) followed by egg-lay 2 (68.4 ± 0.5 %) and egg-lay 3 (57.8 ± 0.5 %; all post-hoc comparisons: P <0.0001, Fig 7B). Although the average number of eggs per female was not affected by gonotrophic cycle (X^2^ = 3.48, P =0.17; Appendix 3-Fig S2A), the average number of larvae varied among gonotrophic cycles (X^2^ = 10.1, P = 0.006), with the highest average in gonotrophic cycle 4 (30 ± 3 larvae per female) followed by gonotrophic cycle 3 (25 ± 3 larvae per female) and gonotrophic cycle 2 (22 ± 1 larvae per female; Tukey’s post-hoc tests significantly different only for the comparison between egg-lay 2 and 4, Appendix 3-Fig S2B). In addition, the survival rate was negatively correlated with the average number of eggs laid (X^2^ = 5.5, P = 0.02; Appendix 4-Supplementary method and results; Appendix 3-Fig S3B).

The parasite isolate had no effect on egg-laying rate (X^2^ = 1.8, P = 0.28). However, females fed blood from isolate A during the first feeding episode laid on average more eggs during gonotrophic cycles 2-4 than females fed blood from isolate B (41±3 and 28±1 eggs respectively; X^2^ = 9.12, P = 0.0025. Appendix 3-Fig S4A), the hatching rate of their eggs was also higher (isolate A: 71.5 ± 0.5 % vs. isolate B: 62.9 ± 0.5 %; X^2^ = 11.58, P <0.001. Appendix 3-Fig S4B) and they produced on average more larvae than females fed on blood from donor B (isolate A: 32 ± 2 and isolate B: 19 ± 1 larvae; X^2^ = 23, P < 0.0001; Appendix 3-Fig S4C).

The blood meal size had no significant effect on egg-laying rate (X^2^ = 0.63, P = 0.43), the average number of eggs (X^2^ = 0.1, P = 0.75), and the average number of 1^st^ instar larvae (X^2^ = 1.09, P = 0.3) but was negatively correlated to the hatching rate (X^2^ = 33.43, P <0.0001).

### F1 development time

Neither the blood type nor the parasite exposure of the mothers significantly influenced the development time of their progeny (X^2^3 = 3.7, P =0.3, Fig 8A and X^2^1 = 0.07, P=0.79, Fig 8B, respectively). The larval density in the rearing cup had no effect on the development time (X^2^1 = 1.47, P=0.22). F1 males developed significantly faster than F1 females (10.65 ± 0.04 days *vs*.10. 52 ± 0.04 days; X^2^1 = 3.98, P = 0.046). The gonotrophic cycle significantly influenced mosquito’s development time (X^2^2 = 64.4, P < 0.0001; Fig 8A). In particular, individuals from gonotrophic cycle 4 developed significantly faster (10.12 ± 0.04 days) than the ones from gonotrophic cycles 2 and 3 (10.67 ± 0.04 days and 10.82 ± 0.05 days, respectively; Tukey’s *post-hoc* tests <0.001, no significant difference between gonotrophic cycles 2 and 3). The sex ratio was not affected by the blood type (X^2^3 = 0.28, P = 0.96).

**Figure 8.**
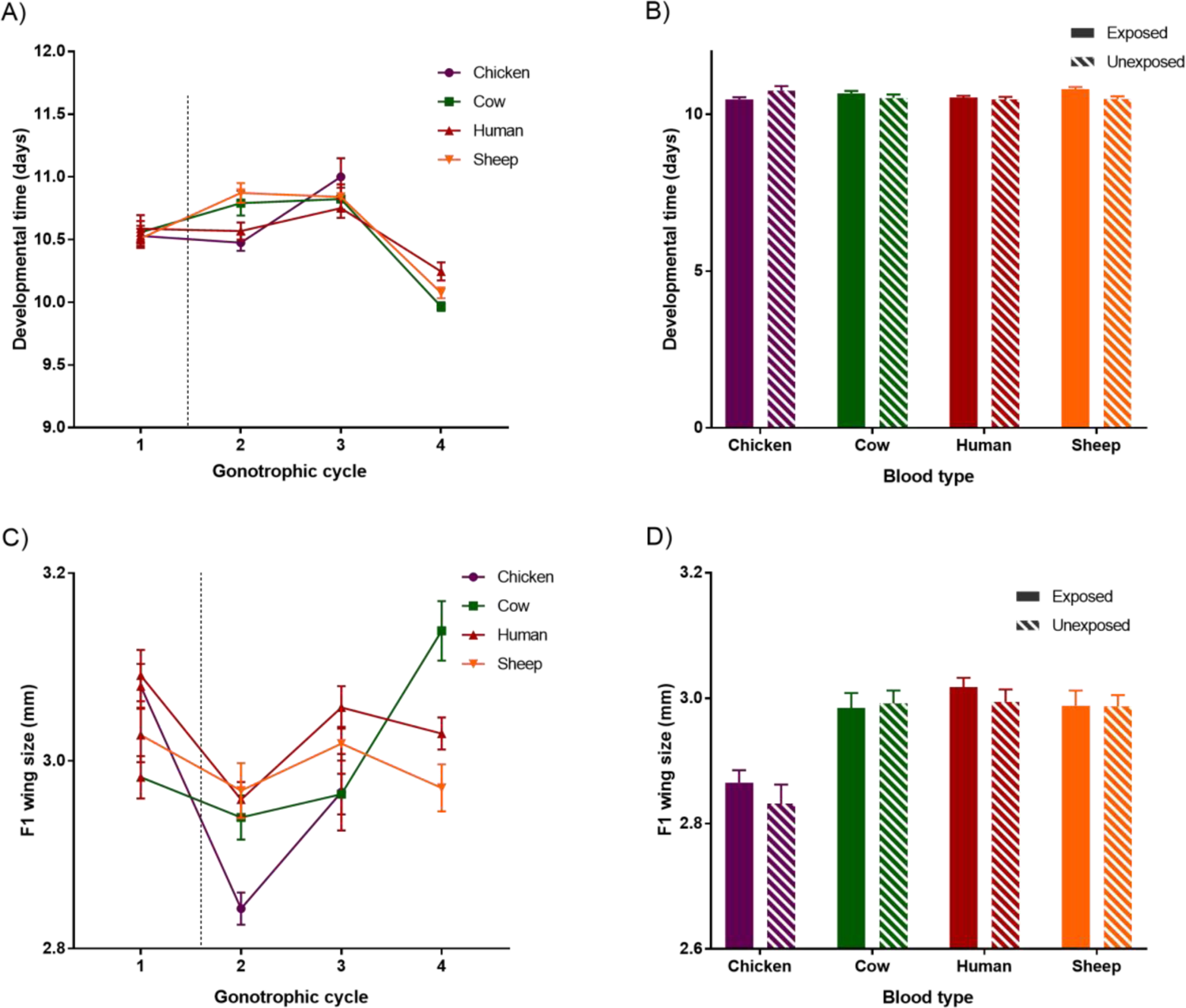
Effects of blood type on progeny development and wing size. A) Effects of blood type on the development time of the progeny for gonotrophic cycles 1, 2, 3 and 4. B) Effects of blood type and infection group (females exposed vs. females unexposed to an infectious blood meal on feeding episode 1) on the development time of the progeny averaged from gonotrophic cycles 2, 3 and 4. C) Effects of blood type on the wing size of the progeny from gonotrophic cycles 1, 2, 3 and 4. D) Effects of blood type and infection group on the average progeny wing size ± se, averaged from gonotrophic cycles 2, 3 and 4.

### F1 wing length

Mosquito wing length was significantly affected by the blood type (X^2^ = 12.9, P = 0.005; Fig 8C). The wing lengths of progeny from females fed on and chicken blood (2.86 ± 0.02mm) were significantly shorter than the progeny from females fed on human blood (3.009 ± 0.01mm; Tukey’s *post-hoc* test: z=3.7, P = 0.001), on cow blood (2.99± 0.02mm; Tukey’s *post-hoc* test: z=2.7, P = 0.04,) and on sheep blood (2.99 ± 0.01mm; Tukey’s *post-hoc* test: z=2.7, P = 0.03; all other comparisons being non-significant). F1 females were significantly bigger than F1 males (3.03 ± 0.01 *vs.* 2.9 ± 0.01mm; X^2^ =132, P<0.0001). Gonotrophic cycle significantly influenced mosquito wing length (X^2^ =30, P<0.0001) with significantly bigger individuals in gonotrophic cycle 3 and 4 compared to gonotrophic cycle 2 (Tukey’s *post-hoc* tests: P<0.001 and P <0.0001, respectively; no significant difference between gonotrophic cycle 3 and 4: Tukey’s *post-hoc* tests: P = 0.26). There was a negative effect of density on F1 size (X^2^ =9.6, P<0.01) and maternal parasite exposure did not significantly affect the progeny wing length (X^2^ = 0.43, P =0.51; Fig 8D).

### Theoretical modelling

Our simulations showed an average IC (the average number of infecting bites transmitted by an infected mosquito during its lifetime) of 0.16 infectious bites when females fed on human hosts only (corresponding to the values with zero feeding attempts on the alternative host in Fig. 9). Compared to a human blood meal, feeding on chicken blood drastically reduced the individual vectorial capacity. There was 2.6 times fewer infectious bites after a single potential blood meal on chicken (with a 0.796 probability of feeding success; mean IC=0.06) and even 26 times fewer after 3 potential blood meals on chicken (mean IC=0.006; Fig. 9). Although less marked, a similar decrease was observed when females obtained blood meals on cow with almost halved of the individual vectorial capacity after 3 potential blood meals on cow (with a 0.717 probability of feeding success; mean IC=0.09; Fig.9). On the contrary, feeding on sheep during *Plasmodium* development increased the vectorial capacity by 25% after 3 potential blood meals on sheep (with a 0.774 probability of feeding success; mean IC=0.2; Fig. 9).

**Figure 9.**
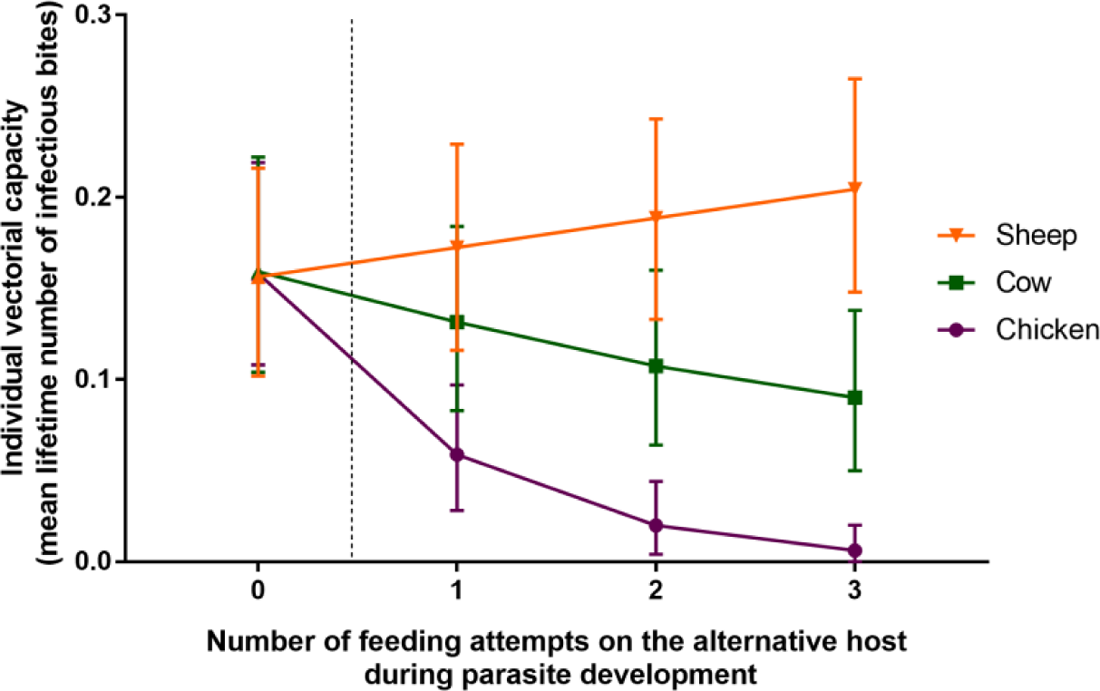
Theoretical modeling of the individual vectorial capacity. (mean lifetime number of infectious bites) at the population level depending on the number of feedings attempted on an alternative host during the parasite development time with zero corresponding to blood meals taken on human host only. The individual vectorial capacity was estimated with a model simulating the daily life history of individual mosquito vectors after taking an infectious blood meal on a human under various scenarios. The scenario was characterized by the presence of humans and an alternative host (either chicken, cow or sheep) with varying availability (0 to 3 consecutive possible feeding attempts during *Plasmodium* incubation period).

## Discussion

The relationships between blood meal type, and mosquito and parasite fitness were explored using a total of 2810 *An. coluzzii* females initially fed an infectious or a non-infectious blood meal and then followed by up to three subsequent blood meals from either human, chicken, cow or sheep. We found no significant effect of blood type on malaria parasite development both at the non-transmissible (oocyst) and the transmissible (sporozoite) stages for either parasite prevalence or intensity. No effects at the oocyst stage or negative effects at the sporozoite stage were also found in *An. gambiae s.s*. females consuming a second blood meal on cow compared to human blood or unfed controls, whereas higher oocyst and sporozoite prevalences were observed in *An. arabiensis* females having a second blood meal on cow compared to human blood or unfed controls (Emami *et al*. 2017). In another study, a second blood meal shortened parasite development in *P. falciparum* with no group specific differences, while a marginal increase was observed with human blood compared to unfed controls for *P. berghei* development (Pathak *et al*. 2022). Thus, the effects of blood type on *Plasmodium* sp. development seem to be variable both between and within *Plasmodium* species and our results with sympatric field strains do not seem to confirm those previous findings. Findings obtained in the laboratory using unnatural host-parasite associations or long-time derived strains do not always reflect natural interactions and an increasing number of studies highlights the importance of confirming laboratory observations with more natural systems for studying disease ecology and evolution (Aguilar *et al*. 2005, Tripet *et al*. 2008).

Exposure to malaria parasite with wild isolates resulted in 78% (isolate A) and 52% (isolate B) infected females and had, overall, no effect on female fitness traits. Parasite exposure did not affect female’s survival regardless of the host type they fed on nor did it affect female fecundity nor their progeny development time and wing size. The only indirect cost we observed was a lower feeding rate in the following blood meals of exposed females compared to unexposed females. The existence of fitness costs of malaria parasite infection in the mosquito host has long been debated and seem to depend on the environmental conditions under which the fitness traits are measured. First, mortality is more commonly reported in unnatural parasite-vector combinations (Ferguson and Read 2002). Second, fitness costs are also more commonly observed in stressful environmental conditions (Lalubin *et al*. 2014, Sangare *et al*. 2014, Roux *et al*. 2015) and can depend on the genetic background (Alout *et al*. 2016). In our experiment, we used a natural parasite-host combination and did not provide any sugar-meals to not alleviate potential fitness costs. Indeed, sugar feeding can affect mosquito competence, survival and fecundity ((Ferguson and Read 2002, Lambrechts *et al*. 2006, Foster 2022) and could compensate for the fitness costs of the different blood types. One explanation could be that the cost of infection are minimal in our system and that infection might only be costly for exposed-infected females. Following the infectious blood meal, our setup did not separate exposed-infected and exposed-uninfected females, thus we were not able to measure the cost of infection in exposed-infected females only. Another possibility is that those costs are quickly offset by the following blood meals the females received, although a study on *Plasmodium relictum* and *Culex pipiens* observed fecundity costs following the infection which lasted for three consecutive gonotrophic cycles (Pigeault and Villa 2018).

Blood type strongly affected mosquito survival. In particular, chicken blood reduced mosquito survivorship by 40% (Fig 6). The larger and nucleated red blood cells of chicken compared to human might be more difficult to digest for *An. coluzzii* (Wintrobe 1933). In addition, although anopheline mosquitoes can reduce their body temperature while blood feeding (Lahondère and Lazzari 2012), chicken host temperature might be too high for evaporative cooling as we saw increased mortality after each blood meal. Interestingly, females fed on sheep blood had the highest survival followed by human which translated in an increased vectorial capacity in our transmission model (Fig 9). Indeed, the type of host the female feeds on strongly increase or decrease the average number of infectious bites in the population (Fig 9). Modeling of malaria transmission showed that mosquito survival is the factor with the biggest impact on transmission (MacDonald 1957). Indeed, parasite development is relatively long (10-14 days) compared to mosquito survival (2-3 weeks). Consequently, females will be infectious for a limited period of time only and any small changes in mosquito longevity will dramatically affect malaria transmission (Smith and McKenzie 2004).

Vectorial capacity is also very sensitive to feeding rates (Brady *et al*. 2016). Our model highlights how even minute differences in survival and feeding rates, such as those observed between females fed with human and sheep blood (Fig 5B & 6), can cause large variation in vectorial capacity (Fig.9). Mosquito feeding rate was highest on sheep blood followed by chicken, human and cow blood. Although membrane feeders were maintained at a temperature corresponding to each vertebrate body temperature, the difference is likely linked to the blood characteristics for the females as sheep blood was maintained at 39°C which is close to cow blood temperature, 38.5°C. Comparison of feeding rates between several vertebrates with blood maintained at the same temperature showed large variation depending on the mosquito species (Phasomkusolsil *et al*. 2013, Al-Rashidi *et al*. 2022). Contrary to our results, sheep blood tended to have a negative impact on survival in *An. dirus*, *An. cracens* and *An. minimus* female and on fecundity in the five mosquito species investigated (Phasomkusolsil *et al*. 2013). Future studies linking fitness measurements and detailed vertebrate blood composition would be useful to determine the chemical and physical characteristics influencing blood digestion and utilization in different mosquito species.

We measured several individual fecundity traits (egg laying rate, average number of eggs, average number of 1^st^ instar larvae, eggs and larvae prevalence, hatching rate) and found no effect of blood type. However, the lifetime fecundity and production of larvae corresponding to the sum of the average number of eggs or larvae of gonotrophic cycles 2, 3 and 4, were affected by the blood type which was lower for chicken compared to human (Fig 7C, D). Thus, our proxy of the lifetime fecundity showed that the fitness of females fed on chicken or cow blood was lower than the fitness of females fed on human or sheep blood. Indeed, the small differences observed at each gonotrophic cycle were not significantly different, but the accumulation of all those small differences gave overall a difference when looking over the lifetime production of eggs and larvae.

We observed a donor effect on fecundity with females fed on blood from donor A on their first blood meal laying more eggs and having a higher hatching rate and more larvae than females fed on donor B. The difference between the two donors in the average number of eggs and larvae in blood meals 2, 3 and 4 tended to blur over the successive gonotrophic cycles (Fig S4A & S4D), such that the successive blood meals slowly offset the difference in donor blood quality.

The survival rate was negatively associated with the average number of eggs laid (Fig S3B) and this association was stronger for the females fed on human blood compared to the other blood types. The life history traits of an organism are constrained by the total amount of resources available (Stearns 1992) such as the tradeoff observed here between the energy allocated to reproduction and survival. Similar results were observed in *Culex pipiens* for which the higher the number of eggs laid, the lower was their subsequent survival (Vézilier *et al*. 2012). Such trade-off between reproduction and survival has been extensively characterized in many organisms with *Drosophila melanogaster* being one of the main model system (Zera and Harshman 2001, Flatt 2011, Flatt 2020, Hsu *et al*. 2021).

Although blood type did not inluence progeny development time, progeny from females fed on chicken blood was smaller than the progeny from females fed on all other vertebrates blood (Fig. 8), which highlights the fitness cost of feeding on chicken blood. Larger females generally take larger blood meals, lay more eggs, live longer and are more competent (Briegel 1990, Kittayapong *et al*. 1992, Lyimo and Koella 1992, Lyimo and Takken 1993, Barreaux *et al*. 2016), but the local environmental conditions can modulate this pattern (Barreaux *et al*. 2016, Barreaux *et al*. 2018). Thus, the bloodmeal type-mediated differences in mosquito progeny size could have strong effects on malaria parasite transmission in the following generations, through increases in mosquito density, competence and survival, three traits that play key role in transmission.

Although we measured several life-history traits simultaneously, our experimental set-up has several limitations. First, we measured averaged fecundity traits per cup since multiple blood-feeding over the lifetime of each female individually would have been technically challenging (i.e. using a single membrane feeder for each individual female), and at the minimum would have strongly reduced our sample size. Second, the laboratory colony used is replenished with wild mosquitoes, however processes such as genetic drift or selection by artificial feeding in the laboratory can happen on a very short time frame especially in small population sizes and those could have eroded the specialization on human blood. Third, specialization on humans could be linked to other ecological or behavioural factors which might exert stronger selection pressures than blood characteristics. While we investigated the effects of the blood type, this was disconnected from the effects of the host type as a whole since other characteristics were not taken into account such as e.g. defensive behavior (Edman and Scott 1987, Lyimo *et al*. 2012), host availability (Lyimo *et al*. 2013, Fikrig *et al*. 2021), parasite manipulation of host behaviour (Vantaux *et al*. 2021) or female individual experience (Vantaux *et al*. 2014). In addition, even though we observed a lower feeding rate of exposed females compared to unexposed females in the following blood meals, all successive blood meals were carried out on membrane feeders and results could be different on whole-body hosts. Fourth, our laboratory setting did not take into account the natural blood foraging rhythm of the vector nor the circadian rhythm of the parasite which have both been shown to influence mosquito and parasite fitness (Schneider *et al*. 2018, O’Donnell *et al*. 2019, Habtewold *et al*. 2022). Fifth, we measured mosquito competence at two time points classically illustrating the non-transmissible and the transmissible parasite stages. It would be interesting to measure the effects of multiple blood meals on different hosts on a more continuous time line as studies showed that multiple blood meals accelerate parasite development (Shaw *et al*. 2021) and growth (Habtewold *et al*. 2021, Kwon *et al*. 2021) and this could have tremendous effects on pathogen development and disease transmission (Brackney *et al*. 2021). Here, successive meals were taken from one of the four vertebrate species used. Under natural conditions, mosquitoes can shift from one host species to another between their gonotrophic cycles. It would be of particular interest to examine the effect of successive meals taken from different host species on the traits measured here. The type and frequency of blood meals not only has consequences on reproduction, survival and epidemiology but also on many other physiological and ecological aspects such as e.g. the maintenance of insecticide resistance phenotype for longer period (Oliver and Brooke 2014, Oliver *et al*. 2022) or an increased ivermectin susceptibility in previously bloodfed females (Seaman *et al*. 2015). Our findings emphasize that considering the diversity of vertebrate blood-meal sources is important to better understand the ecology of mosquitoes as well as their capacity to transmit malaria parasites.

Overall, blood type had a significant impact on mosquito survival and feeding rates, leading to considerable variation in vectorial capacity and differences in progeny sizes. These findings imply that the diversity of vertebrate hosts (including both the number of species and their relative abundance) within villages could influence the transmission of malaria parasites. Specifically, transmission may decrease when chickens or cows make up the majority of available blood sources, while it may increase when a relatively large number of sheep are present. However, the host selection patterns of mosquitoes are not solely driven by vertebrate abundance, but are also influenced by mosquito innate preferences and host defensive behavior (Lefèvre *et al*. 2009, Lyimo and Ferguson 2009). The human blood index (number of females fed on humans, including mixed human-animal bood meals, over the total number of blood fed females) was highly variable between villages and mosquito species in this area of Burkina Faso, with anthropophagy ranging from 56.5± 4% to 83.5 ± 2.2% in different villages (Vantaux *et al*. 2021). Among the 2627 fed *An. gambiae s.l*. for which the blood meal origin was determined during this study, only 10 (0.38± 0.24%) individuals fed on chicken, 63 (2.4± 0.6%) individuals fed on sheep and 951 (36.2± 1.8%) fed on cow (A. Vantaux personnal communication). Although those numbers represent only a limited period of the year, they are a reminder that *An. gambiae s.l*. females also feed on non-human hosts and that the local composition of domestic animals could influence parasite transmission in the village. Limiting or increasing access to particular non-human hosts in the village could help improve malaria control during the peak of transmission. Since zooprophylaxis, which involves using livestock to draw mosquitoes away from humans and decrease malaria transmission, is dependant on specific conditions such as the optimal distance between humans and livestock sleeping areas and the characteristics of the local mosquito populations (Donnelly *et al*. 2015, Hasyim *et al*. 2018), our findings suggest that it may be feasible to further potentiate zooprophylaxis effects by considering the use of endectocides (Pooda *et al*. 2015, Khaligh *et al*. 2021) and/or application of formulation on animal fur to divert mosquitoes from their prefered host to less prefered hosts (Kemibala *et al*. 2020).

## Acknowledgements

We would like to thank all volunteers for participating in this study as well as the local authorities for their support. We are very grateful to the IRSS staff in Burkina Faso for technical assistance. We thank Diego Santiago-Alarcon, Francisco C. Ferreira and an anonymous reviewer for their fruitful comments.

## Data, scripts, code, and supplementary information availability

Data and script for the statistical analyses are available online: https://zenodo.org/record/7940843

Data and scripts for the theoretical modeling are available online: https://zenodo.org/record/7645483

