## supplementary files for "Multiple hosts, multiple impacts: the role of vertebrate host diversity in shaping mosquito life history and pathogen transmission"

### Appendix

Appendix 1 - Table S1: Parasite prevalence and intensity in females.

| Replicate | Parasite isolate | Gametocyte Density (µl) | Blood meal type | Prevalence ± 95% CI | Parasite/midgut ± se | n |
|---|---|---|---|---|---|---|
| 1 | A | 232 | Chicken | 1 ± 0 | 50.5 ± 15.5 | 2 |
| | | | Cow | 0.43 ± 0.37 | 8.3 ± 4.7 | 7 |
| | | | Human | 0.85 ± 0.2 | 15 ± 3.9 | 13 |
| | | | Sheep | 0.86 ± 0.18 | 25 ± 6.1 | 14 |
| 2 | B | 32 | Chicken | 0.7 ± 0.24 | 7.9 ± 1.6 | 15 |
| | | | Cow | 0.43 ± 0.26 | 4.2 ± 1 | 14 |
| | | | Human | 0.45 ± 0.21 | 4.7 ± 0.8 | 22 |
| | | | Sheep | 0.53 ± 0.17 | 8.2 ± 1.2 | 32 |
| 3 | C | 136 | Chicken | 0.56 ± 0.13 | 8.6 ± 1.2 | 52 |
| | | | Cow | 0.65 ± 0.13 | 10 ± 1.4 | 49 |
| | | | Human | 0.87 ± 0.09 | 7.2 ± 0.9 | 52 |
| | | | Sheep | 0.9 ± 0.08 | 6.8 ± 0.9 | 49 |
| 4 | D | 72 | Chicken | 0.76 ± 0.17 | 8.4 ± 1.3 | 25 |
| | | | Cow | 0.78 ± 0.11 | 11.7 ± 1.2 | 54 |
| | | | Human | 0.76 ± 0.1 | 6.4 ± 0.8 | 66 |
| | | | Sheep | 0.73 ± 0.1 | 8.5 ± 1 | 70 |
| 5 | E | 192 | Chicken | 0.84 ± 0.14 | 33.5 ± 5.3 | 25 |
| | | | Cow | 0.81 ± 0.11 | 21.2 ± 3.1 | 47 |
| | | | Human | 0.77 ± 0.12 | 26.6 ± 2.9 | 47 |
| | | | Sheep | 0.84 ± 0.1 | 24.1 ± 2.7 | 49 |
| | F | 136 | Chicken | 0.94 ± 0.08 | 15.6 ± 2.3 | 32 |
| | | | Cow | 0.71 ± 0.13 | 19.6 ± 2.5 | 51 |
| | | | Human | 0.74 ± 0.12 | 18.4 ± 2.5 | 47 |
| | | | Sheep | 0.74 ± 0.13 | 19.7 ± 2.6 | 42 |

Appendix 2 - Table S2- Statistical analyses results

| Life-history traits | Response variable | Model | Effect Test Outputs |
|---|---|---|---|
| Competence | Oocyst prevalence, design 1 | GLMM, binomial distribution | Blood type: LRT $X^2_3$ = 5.02, P = 0.17<br>**Parasite isolate: LRT $X^2_1$ = 6.92, P = 0.009** |
| | Oocyst intensity, design 1 | GLMM, zero-truncated negative binomial distribution | **Blood type: LRT $X^2_3$ = 10.55, P = 0.01**<br>**Parasite isolate: LRT $X^2_1$ = 24.89, P < 0.0001** |
| | Sporozoite prevalence, design 1 | GLMM, binomial distribution | Blood type: LRT $X^2_2$ = 0.62, P = 0.73<br>Parasite isolate: LRT $X^2_1$ = 2.09, P = 0.15 |
| | Oocyst prevalence, design 2 | GLMM, binomial distribution | Blood type: LRT $X^2_3$ = 3.14, P = 0.37<br>Gametocytemia: LRT $X^2_1$= 1.37, P = 0.24<br>Blood type*Gametocytemia: LRT $X^2_3$= 1.27, P = 0.74 |
| | Oocyst intensity, design 2 | GLMM, zero-truncated negative binomial distribution | Blood type: LRT $X^2_3$= 4.99, P = 0.17<br>**Gametocytemia: LRT $X^2_1$=11.09, P =< 0.001**<br>**Blood type*Gametocytemia: LRT $X^2_3$= 11.06, P = 0.01** |
| | Sporozoite prevalence, design 2 | GLMM, binomial distribution | Blood type: LRT $X^2_2$ = 0.65, P = 0.72<br>Gametocytemia: LRT $X^2_1$ = 1.3, P = 0.25<br>Blood type*Gametocytemia: LRT $X^2_2$ = 0.33, P = 0.85 |
| Feeding rate | Proportion of fed females during blood meal 2 to 4 | GLMM, binomial distribution | **Blood type: LRT $X^2_3$ = 14.4, P = 0.002**<br>**Feeding episode: LRT $X^2_2$ = 160.2, P< 0.0001**<br>**Exposure: LRT $X^2_1$ = 9.09, P = 0.003**<br>Parasite isolate: LRT $X^2_1$ = 1.17, P = 0.28<br>**Blood type*Feeding episode: LRT $X^2_6$ = 37.8, P< 0.0001**<br>Blood type*Exposure: LRT $X^2_3$ = 0.38, P=0.94<br>Exposure*Feeding episode: LRT $X^2_2$ = 0.03, P=0.99<br>Exposure*Feeding episode*Blood type: LRT $X^2_6$ = 12.29, P=0.06 |
| Mosquito blood meal size | Size blood meal 1 | ANOVA | **Parasite isolate: $F_{1, 115}$=34.6, P < 0.0001**<br>Exposure: $F_{1, 113}$=0.48, P = 0.49<br>Isolate*Exposure: $F_{1, 113}$=0.14, P = 0.71 |
| | Size blood meal 2 to 4 | GLMM, Gaussian distribution | Blood type: LRT: $X^2_3$ = 4.2, P = 0.24<br>**Feeding episode: LRT $X^2_2$ = 54.04, P < 0.0001**<br>Exposure: LRT $X^2_1$ = 0.18, P = 0.67<br>Parasite isolate: LRT $X^2_1$ = 2.4, P = 0.12<br>**Blood type*Feeding episode: LRT $X^2_6$ = 23.7, P = 0.0006**<br>Blood type* Exposure: LRT $X^2_3$ = 4.7, P = 0.19<br>Feeding episode*Exposure: LRT $X^2_2$ = 0.8, P = 0.67<br>Exposure*Feeding episode*Blood type: LRT $X^2_6$ = 2.5, P = 0.87 |
| Survival | Survival | Cox proportional hazard mixed models | **Blood type : LRT $X^2_3$ = 68.26, P < 0.0001**<br>**Parasite isolate: LRT $X^2_1$ = 24.38, P < 0.0001**<br>Exposure : LRT $X^2_1$ =0.03, P=0.87<br>Blood type*Exposure: LRT $X^2_3$=2.38, P=0.50<br>Parasite isolate*Exposure: LRT $X^2_1$=1.25, P=0.26<br>Blood type*Parasite isolate : LRT $X^2_3$ =0.27, P=0.96<br>Blood type*Parasite isolate*Exposure : LRT $X^2_3$=4.79, P =0.19 |
| Fecundity | Egg-laying rate gonotrophic cycle 1 | GLM, binomial distribution | Parasite isolate: LRT $X^2_1$ = 3.2, P = 0.07<br>Exposure : LRT $X^2_1$ = 3.07, P = 0.08<br>Meal size : LRT $X^2_1$ = 0.73, P = 0.39<br>Parasite isolate*Exposure: $X^2_1$ = 1.3E-8, P = 0.99 |
| | Egg-laying rate gonotrophic cycle 2 to 4 | GLMM, binomial distribution | Blood type: LRT $X^2_3$ = 5.6, P = 0.13<br>Meal size: LRT $X^2_1$ = 0.63, P = 0.43 |

| | | | |
|---|---|---|---|
| | | | **Gonotrophic cycle: LRT $X^2_2$ = 53.6, P < 0.0001**<br>Exposure: LRT $X^2_1$ = 0.08, P = 0.78<br>Parasite isolate: LRT $X^2_1$ = 1.8, P = 0.28 |
| | Hatching rate gonotrophic cycle 1 | GLM, quasi binomial distribution | Exposure: LRT $F_{1, 102}$= 0.99, P = 0.32<br>Meal size: LRT $F_{1, 100}$= 0.84, P = 0.36<br>**Parasite isolate : LRT $F_{1, 102}$=67.59, P < 0.0001**<br>Exposure*Parasite isolate: LRT $F_{1, 100}$=0.76, P = 0.38 |
| | Hatching rate gonotrophic cycle 2 to 4 | GLMM, binomial distribution | Blood type: LRT $X^2_3$ = 5.26, P = 0.15<br>Exposure: LRT $X^2_1$ = 2.06, P = 0.15<br>Meal Size : $X^2_1$ = 33.43**, P < 0.0001**<br>**Gonotrophic cycle: LRT $X^2_2$ = 501, P < 0.0001**<br>**Parasite isolate: LRT $X^2_1$ = 11.58, P <0.001** |
| | Average number of eggs gonotrophic cycle 1 | GLM, quasipoisson distribution | Exposure: LRT $F_{1, 106}$= 0.22, P = 0.85<br>**Meal size: LRT $F_{1, 108}$= 6.29, P = 0.014**<br>**Parasite isolate: LRT $F_{1, 108}$=41.54, P < 0.0001**<br>Parasite isolate*Exposure: LRT $F_{1, 106}$= 0.01, P = 0.92 |
| | Average number of eggs gonotrophic cycle 2 to 4 | GLMM, Gaussian distribution | Blood type: LRT $X^2_3$ = 0.85, P = 0.84<br>Meal size: LRT $X^2_1$ = 0.1, P = 0.75<br>Gonotrophic cycle: LRT $X^2_2$ = 3.48, P =0.17<br>Exposure: LRT $X^2_1$ = 0.0005, P = 0.98<br>**Parasite isolate: LRT $X^2_1$ = 9.12, P = 0.0025** |
| | Average number of 1st instar larvae gonotrophic cycle 1 | GLM, quasipoisson distribution | Exposure: LRT $F_{1,100}$=0.02, P = 0.88<br>**Meal size : LRT $F_{1,102}$=4.46, P = 0.04**<br>**Parasite isolate: LRT $F_{1,102}$=56.98, P < 0.0001**<br>Parasite isolate* Exposure: LRT $F_{1, 100}$=0.09, P=0.76 |
| | Average number of 1st instar larvae gonotrophic cycle 2 to 4 | GLMM, Gaussian distribution | Blood type : LRT $X^2_3$ = 1.4, P = 0.70<br>Exposure: LRT $X^2_1$ = 0.97, P = 0.32<br>Meal size: LRT $X^2_1$ = 1.09, P = 0.30<br>**Gonotrophic cycle: LRT $X^2_2$ = 10.1, P = 0.006**<br>**Parasite isolate: LRT $X^2_1$ = 23, P < 0.0001** |
| | Average lifetime fecundity of eggs over gonotrophic cycle 2-4 | GLM, quasipoisson distribution | **Blood type: LRT $F_{1,88}$=4.79, P=0.004**<br>Parasite isolate: LRT $F_{1,84}$=0.02, P=0.90<br>Exposure: LRT $F_{1,84}$=0.005, P=0.94<br>Blood type*Parasite isolate: LRT $F_{1,76}$=0.58, P=0.63<br>Blood type*Exposure: LRT $F_{1,80}$=0.67, P=0.57<br>Exposure*Parasite isolate : LRT $F_{1,80}$=0.06, P=0.81<br>Blood type*Exposure*Parasite isolate : LRT $F_{1,76}$=1.11, P=0.35 |
| | Average lifetime fecundity of 1st instar larvae over gonotrophic cycle 2-4 | GLM, quasipoisson distribution | **Blood type: LRT $F_{1,88}$=3.86, P=0.012**<br>Parasite isolate: LRT $F_{1,84}$=0.63, P=0.43<br>Exposure: $F_{1,84}$=0.07, P=0.79<br>Blood type*Parasite isolate: $F_{1,80}$=0.31, P=0.82<br>Blood type*Exposure: LRT $F_{1,80}$=0.86, P=0.46<br>Exposure*Parasite isolate: LRT $F_{1,76}$=0.0059, P=0.94<br>Blood type*Exposure*Parasite isolate: LRT $F_{1,76}$=0.9, P=0.44 |
| Development time | Development time of larvae from gonotrophic cycle 1 | Cox proportional hazard mixed models | Mosquito sex: $X^2_1$ = 1.23, P = 0.27<br>Maternal parasite exposure: $X^2_1$ = 0.15, P = 0.7<br>Maternal parasite exposure*Mosquito sex : $X^2_1$ = 0.5, P = 0.48 |
| | Development time of larvae from gonotrophic cycle 2-4 | Cox proportional hazard mixed models | Blood type: $X^2_3$ = 3.7, P =0.3<br>Density: $X^2_1$ = 1.5, P =0.23<br>Maternal parasite exposure: $X^2_1$ = 0.07, P=0.79<br>**Mosquito sex : $X^2_1$ = 3.98, P = 0.046**<br>**Gonotrophic cycle : $X^2_2$ = 64.4, P < 0.0001** |

| | | | |
|---|---|---|---|
| F1 wing length | F1 wing length gonotrophic cycle 1 | GLMM, Gaussian distribution | M**osquito sex: $X^2_1$=68, P<0.0001**<br>Maternal parasite exposure: $X^2_1$ = 1.1, P = 0.29<br>Maternal parasite exposure*Mosquito sex : $X^2_1$ = 0.24, P =0.62 |
| | F1 wing length gonotrophic cycle 2-4 | GLMM, Gaussian distribution | **Blood type: $X^2_3$ = 12.9, P = 0.005**<br>**Mosquito sex: $X^2_1$=132, P<0.0001**<br>**Gonotrophic cycle : $X^2_2$=30, P<0.0001**<br>**Density: $X^2_1$=9.64, P=0.0019**<br>Maternal parasite exposure $X^2_1$ = 0.43, P =0.51 |

Appendix 3 – Supplementary Figures

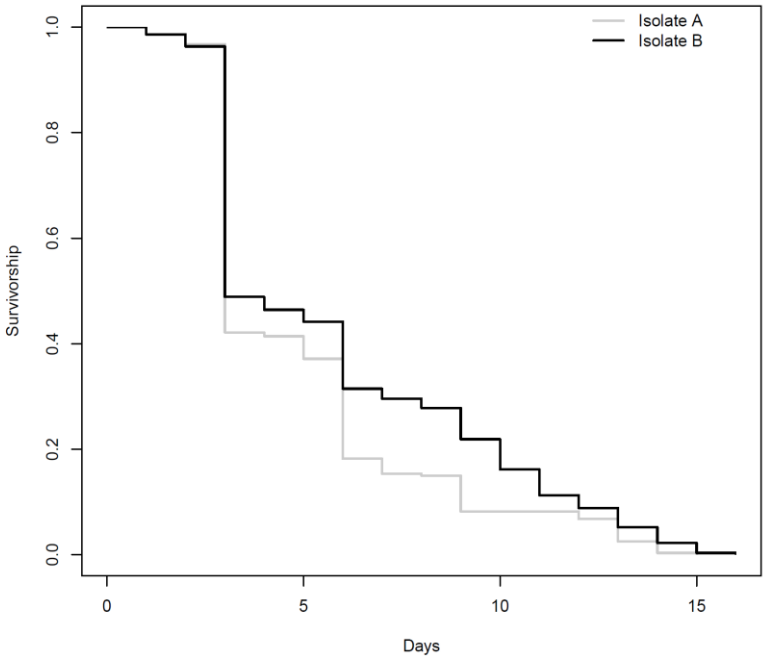


Figure S1. Effects of isolates used for the first infectious blood meal on mosquito survivorship.

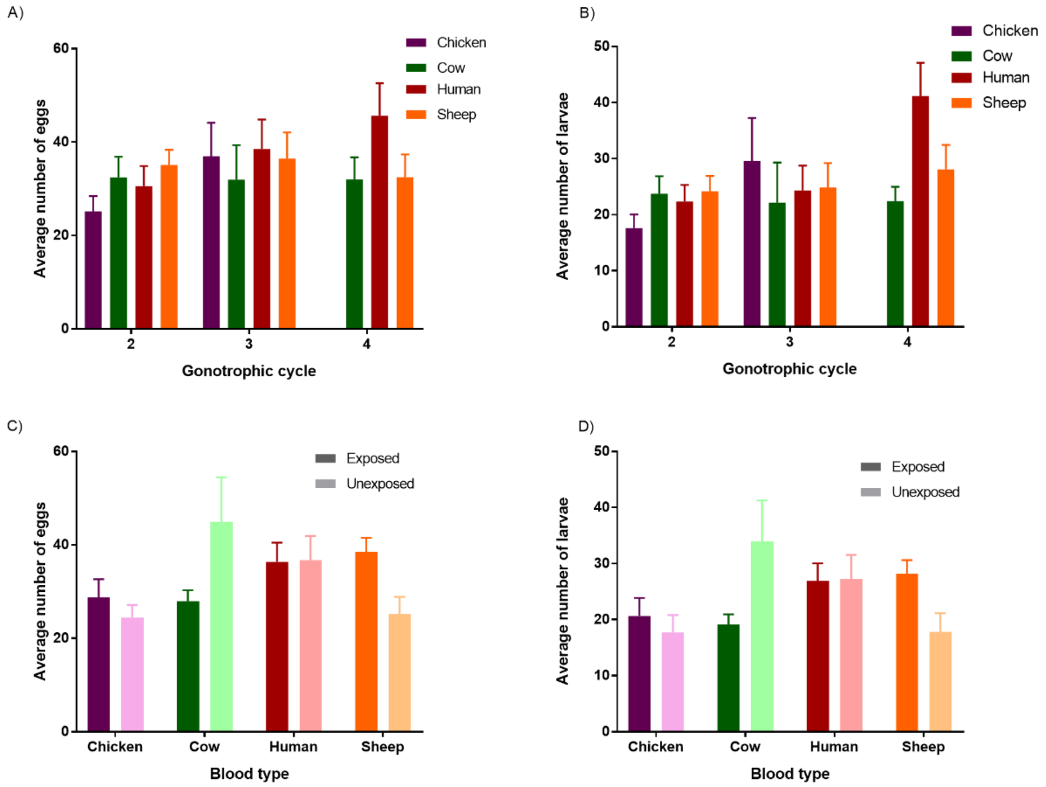


Figure S2. Effects of blood type on A) the average number of eggs per female ± se and B) the average number of larvae per female ± se, from gonotrophic cycles 2, 3 and 4. Effects of blood type and infection group (females exposed vs. females unexposed to an infectious blood meal on feeding episode 1) on C) the average number of eggs per female ± se and on D) the average number of larvae per female ± se averaged from gonotrophic cycles 2, 3 and 4.

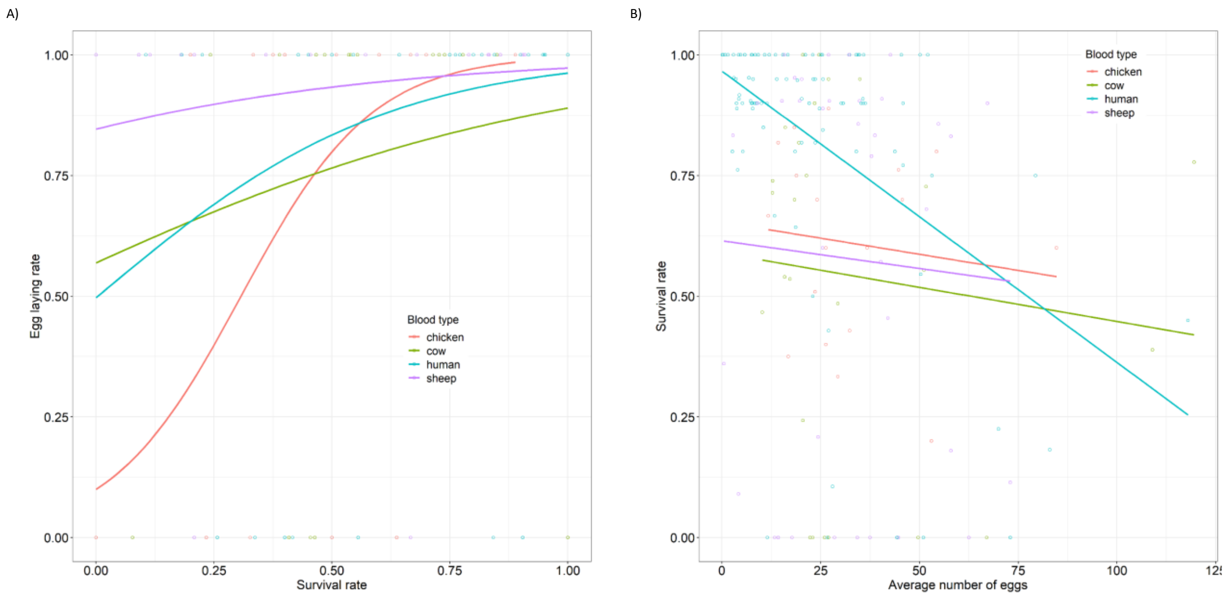


Figure S3. Effect of blood type on A) the relationship between egg laying rate and mosquito survivorship and on B) the relationship between mosquito survivorship and the average number of eggs.

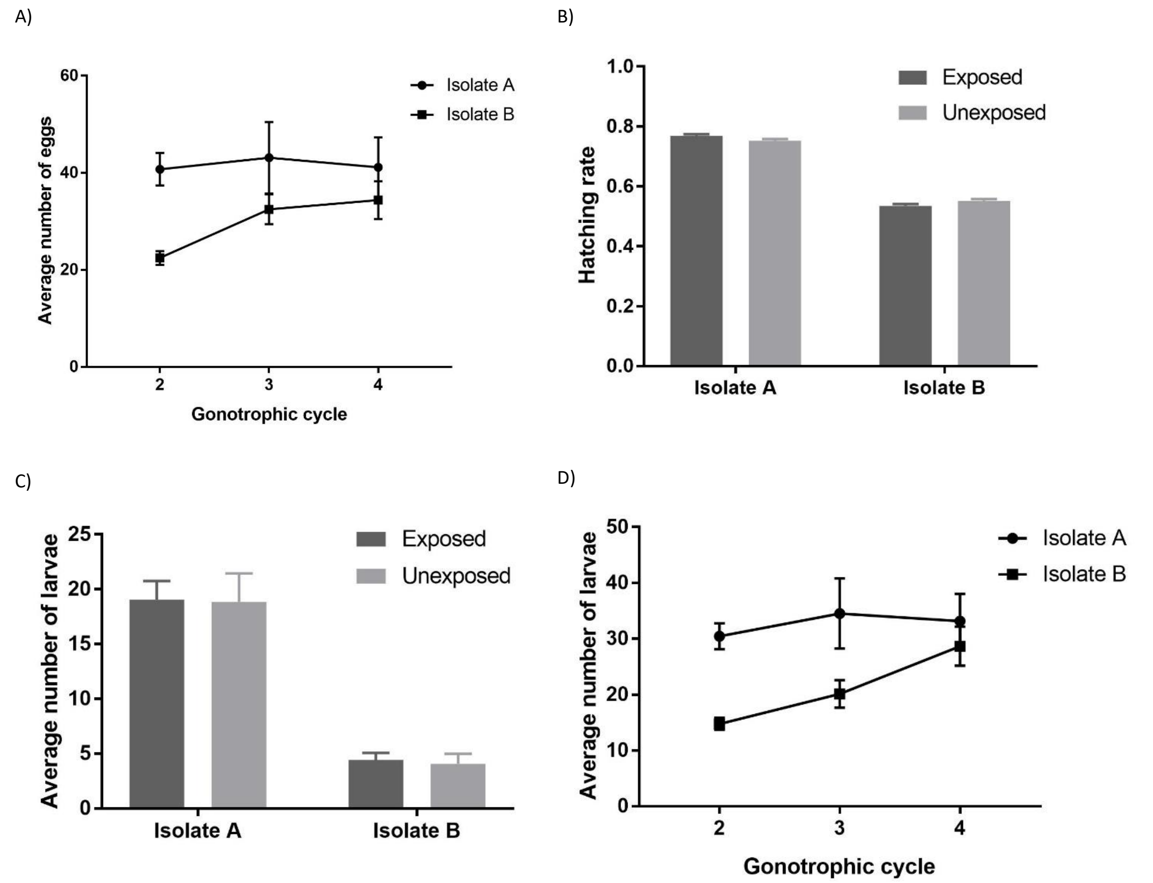


Figure S4. A) Effects of isolates used for the first infectious blood meal on the average number of eggs per female ± se for gonotrophic cycles 2, 3 and 4. B) Effects of isolates used for the first infectious blood meal and infection group (females exposed vs. females unexposed to an infectious blood meal on feeding episode 1) on the hatching rate (number of larvae/number of eggs) ± 95% CI averaged from gonotrophic cycles 2, 3 and 4. C) Effects of isolates used for the first infectious blood meal and infection group (females exposed vs. females unexposed to an infectious blood meal on feeding episode 1) on the average number of larvae per female ± se averaged from gonotrophic cycles 2, 3 and 4. D) Effects of isolates used for the first infectious blood meal on the average number of eggs per female ± se for gonotrophic cycles 2, 3 and 4.

Appendix 4 – Supplementary method and results

**Method**

*Mosquito blood meal size* – Data from the first blood meal were reduced centered (aka centered and scaled) and analyzed using an ANOVA with parasite isolate, mosquito exposure and their interaction as factors.

*Fecundity* – The effects of parasite exposure, isolate and their interaction on egg laying rate , hatching rate, average number of eggs, and average number of 1$^{st}$ instar larvae at the first gonotrophic cycle were analysed using General Linear Models (GLMs) with binomial, quasi binomial, and quasipoisson error structure, respectively.

The effects of blood type, survival rate and their interaction on the egg-laying rate over the four gonotrophic cycles were analyzed with a binomial GLMM with rearing cup as a random factor.

The effects of blood type, the average number of eggs and their interaction on the survival rate over the four gonotrophic cycles were analyzed with a binomial GLMM with rearing cup as a random factor.

The *development time* of larvae from the first gonotrophic cycle was analysed using a Cox proportional hazard mixed effect model with maternal exposure, sex and their interaction coded as fixed factors, and rearing plastic cup as a random factor.

*F1 wing length*–Wing length of the progeny from gonotrophic cycle 1was analysed using a Gaussian GLMM with maternal exposure, mosquito sex and their interactions as fixed factors and rearing cup as a random factor.

**Results**

*Mosquito blood meal size* – Following the first feeding episode on human blood, females exposed to parasite isolate A had smaller blood meals than the ones fed on isolate B (2.21 ± 0.13 µg *vs*. 3.25 ± 0.12 µg respectively; $F_{1, 115}$=34.6, P < 0.0001, Fig S5). The meal size of females fed on infectious blood was similar to that of females fed a noninfectious blood (2.76 ± 0.12µg *vs*. 2.61± 0.19 µg respectively; $F_{1, 113}$=0.37, P = 0.49, Fig S5), regardless of the isolate (parasite isolate*infection status: $F_{1, 113}$=0.14, P = 0.71, Fig S5).

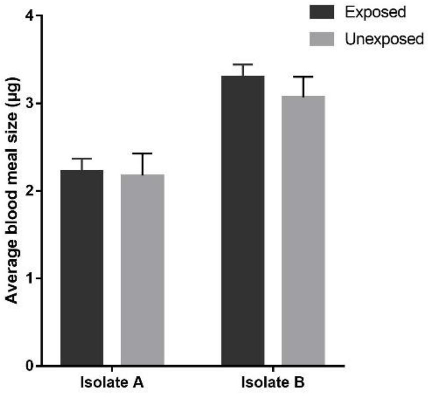


Figure S5. Effects of exposure and parasite isolate on the average blood meal size ± se of the first feeding episode.

*Fecundity* – The egg-laying rate following the first gonotrophic cycle was high with eggs present in 95 ± 4% (Fig 7A) of all cups (n=121). There was no effect of blood meal size ($X^2_1$ = 0.73, P = 0.39), of isolate ($X^2_1$ = 3.2, P = 0.07), nor of parasite exposure ($X^2_1$ = 3.07, P = 0.08) nor of isolate by parasite exposure interaction ($X^2_1$ = 1.3E-8, P = 0.99) on egg-laying rate.

The average number of eggs laid on the first gonotrophic cycle was 16 ± 1 eggs per female (min = 0.1 eggs per females and max=52.1 eggs per female). Females exposed to an infectious blood meal on the first gonotrophic cycle laid as many eggs as unexposed control females (16 ± 1 and 16 ± 3 eggs respectively; $F_{1, 106}$= 0.99, P = 0.32; Fig S6). The average number of eggs laid was negatively correlated to the average blood meal size ($F_{1, 108}$= 6.29, P = 0.014). There was a strong effect of isolate, with females fed blood from isolate A laying more eggs than females fed blood from isolate B (24 ± 2 and 7 ± 1 eggs respectively; $F_{1, 102}$= 67.59, P <0.0001), regardless of parasite exposure (exposure* parasite isolate: $F_{1, 106}$= 0.01, P = 0.92; FigS6).

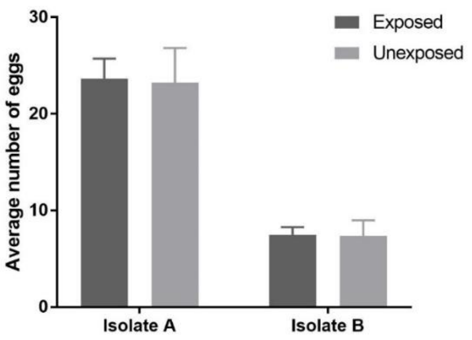


Figure S6. Effects of exposure and parasite isolate on the average number of eggs ± se of the first feeding episode.

On the first gonotrophic cycle 68.1 ± 0.6% of the eggs hatched. Eggs from females exposed to an infectious blood meal had a similar hatching rate as eggs from unexposed females (71 ± 0. 6 % and 68 ± 0.6 % respectively; $F_{1,\ 102}$= 0.99, P = 0.32, Fig S7). Eggs from females fed on isolate A had a higher hatching rate than eggs from females fed on isolate B (76.4 ± 0.6 % and 53.9 ± 0.7 % respectively; $F_{1,\ 102}$=67.59, P < 0.0001), regardless of parasite exposure (exposure* parasite isolate: $F_{1,\ 100}$=0.76, P = 0.38, Fig S7). There was no effect of the average blood meal size ($F_{1,\ 108}$= 6.29, P = 0.014).

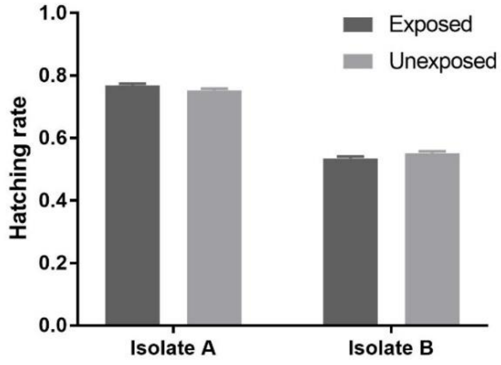


Figure S7. Effects of exposure and parasite isolate on the hatching rate ± 95% CI of the first feeding episode.

The average number of 1$^{st}$ instar larvae on the first gonotrophic cycle was 12 ± 1 larvae per female. There was no effect of parasite exposure on the average number of larvae ($F_{1,100}$=0.02, P = 0.88, Fig S8). Females fed on isolate A had significantly more larvae per female than females fed on isolate B (19 ± 1 and 4 ± 1 larvae per female respectively; $F_{1,102}$=56.98, P < 0.0001), regardless of parasite exposure (exposure*parasite isolate $F_{1,\ 100}$=0.09, P = 0.76, Fig S8). The average number of 1$^{st}$ instar larvae laid was negatively correlated to the average blood meal size ($F_{1,\ 108}$= 6.29, P = 0.014).

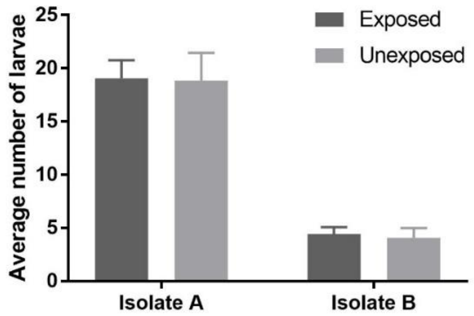

Figure S8. Effects of exposure and parasite isolate on the average number of larvae ± se of the first feeding episode.

The egg-laying rate decreased with the survival rate ($X^2_1 = 32.9$, $P < 0.0001$; Fig S3A). Neither the blood type ($X^2_3 = 7.7$, $P = 0.052$) nor the two-way interaction ($X^2_3 = 7.6$, $P = 0.054$) significantly affected the egg-laying rate.

The survival rate was negatively associated with the average number of eggs laid ($X^2_1 = 5.5$, $P = 0.02$; Fig S3B) and this association was stronger for the females fed on human blood compared to the other blood types ($X^2_3 = 10.2$, $P = 0.017$; Fig S3B). No effect of the interaction was found ($X^2_3 = 3.45$, $P = 0.33$).

*F1 development time* –the F1 development time on the first gonotrophic cycle (10.55±0.04 days) was not affected by mosquito sex ($X^2_1 = 1.23$, $P = 0.27$), nor by maternal parasite exposure ($X^2_1 = 0.15$, $P = 0.7$), nor by their interaction ($X^2_1 = 0.5$, $P = 0.48$).

*F1 wing length* – On the first gonotrophic cycle, F1 females were significantly bigger than F1 males (3.14 ± 0.02 *vs.* 2.97 ± 0.02mm; $X^2_1=68$, $P<0.0001$). Neither maternal parasite exposure ($X^2_1 = 1.1$, $P = 0.29$) nor the two-way interaction ($X^2_1 = 0.24$, $P =0.62$) significantly affected the progeny wing length of gonotrophic cycle 1.
